# Resolving Cargo-motor-track Interactions in Living Cells with Bifocal Parallax Single Particle Tracking

**DOI:** 10.1101/2020.07.06.189746

**Authors:** Xiaodong Cheng, Kuangcai Chen, Bin Dong, Seth L. Filbrun, Gufeng Wang, Ning Fang

## Abstract

Resolving coordinated biomolecular interactions in living cellular environments is vital for understanding the mechanisms of molecular nanomachines. The conventional approach relies on localizing and tracking target biomolecules and/or subcellular organelles labeled with imaging probes. However, it is challenging to gain information on rotational dynamics, which can be more indicative of the work done by molecular motors and their dynamic binding status. Herein, a bifocal parallax single particle tracking method using half-plane point spread functions has been developed to resolve the full-range azimuth angle (0-360°), polar angle, and 3D displacement in real time under complex living cell conditions. Using this method, quantitative rotational and translational motion of the cargo in a 3D cell cytoskeleton was obtained. Not only well-known active intracellular transport and free diffusion were observed but new interactions, tight attachment and tethered rotation, were discovered for better interpretation of the dynamics of cargo-motor-track interactions at various types of microtubule intersections.

**Significance Statement:** Translation and rotational motion of cargo during pauses at the microtubule intersections in living cells were revealed by high-accuracy three-dimensional single particle rotational tracking. The current study demonstrates the potential of studying coordinated interactions in living cellular environments by resolving characteristic rotational motions.

## Introduction

Coordinated biomolecular interactions are essential to molecular nanomachines in fulfilling their functions. The related structural and dynamics information is crucial in elucidating the underlying working mechanisms of molecular nanomachines. Biochemical assays, crystallography, cryo-electron microscopy and force spectroscopy are well-developed tools to study binding affinity, structure and force (1-4). On the other hand, single particle tracking (SPT) has become an effective approach to reveal the dynamics of molecular interactions in living cells, including the timing, location, velocity, diffusion, local viscosity, etc., from trajectories of individual imaging probes (5-9). The information derived from localization is mostly about the *translational* motion; however, it does not reveal the *rotational* motion, which is either induced directly by molecular motors, such as ATP synthase (10) and dynamin helices (11), or a reflection of dynamic binding status, e.g., tight or loose binding, single or multiple attachment points. There is an imperative need for reliable measurements of the rotational motion in living cells in order to complete our interpretation of the dynamics of molecular interactions.

A fundamental principle of tracking the rotational motion is to resolve the orientation (azimuth and polar angles) of anisotropic imaging probes, such as fluorescent dipolar emitters and plasmonic gold nanorods (GNRs), in real time using either polarization-based (12-18) or defocused (19-22) imaging techniques. Despite the previous successes, however, several persistent challenges remain for three-dimensional (3D) single particle rotational tracking in living cells. Firstly, tracking rotational probes in the presence of movement along the axial direction (z-axis) is challenging because the signal changes induced by the axial movement can be easily confused with those generated by rotation. Secondly, angular degeneracy is a common limitation of the polarization-based techniques. The resolvable angle range with one set of orthogonal polarization directions is limited to 0-90°, and further polarization splitting or defocusing is needed to expand the angle range, which, however, inevitably makes negative impacts on 3D localization and angle estimation. The analysis of the electric field distribution in the forms of defocused image patterns is currently the best method to circumvent the angular degeneracy issue and discover the dipole’s orientation in the full angle range; however, it has never been realized for 3D dynamic tracking due to inconsistent image patterns as the defocusing distance changes continuously with the target probe’s movement along the z-axis.

In this study, we report a bifocal parallax SPT technique to overcome the aforementioned challenges and realize the highest-to-date accuracy in simultaneous localization and orientation determination of anisotropic imaging probes in living cells by combining bifocal (one focused channel and one defocused channel with a fixed defocusing distance) dark field microscopy with a parallax imaging and auto-focusing system. Using the newly-established ability to dynamically reveal the rich information of anisotropic imaging probes, we visualized and interpreted the dynamics of cargo-motor-track interactions during the cargo’s intracellular transport crossing the 3D microtubule network in living cells with unprecedented details. Understanding the dynamics of binding events between the cargo, motor and microtubule provides a key perspective to decipher how the motor proteins compete and coordinate to transport the cargo on the 3D microtubule geometry (23-27). Microtubule intersections play a significant role in regulating the cargo’s moving direction towards various destinations inside the cell (28, 29). Studies of the cargo’s transport dynamics (30, 31), force generated by the molecular motors (32, 33), and effects of 3D microtubule geometry (34, 35) have provided insights on the working mechanisms that govern the cargo behaviors at the track intersections. The current effort is dedicated to expanding the understanding of the cargo behaviors by uncovering the correlation between the cargo’s rotational motion and the cargo-motor-microtubule binding dynamics, as well as the torques exerted by molecular motors, in living cells.

## Materials and Methods

### Preparation of transferrin@GNRs

Gold nanorods stabilized by sodium citrate were obtained from Nanopartz (Salt Lake City, UT). The average aspect ratio of the GNRs was 2.0. The electrostatic adsorption method establishing transferrin-coated GNRs was reported previously in the literature (36). 4 µL of 50 mM borate buffer (pH 8.5) and 5 µL of 3mg/mL transferrin (CAS: 11096-37-0, Sigma-Aldrich) was mixed with 100 µL GNRs solution (3.7×10^10^ particles/mL). The mixture was left to rest at room temperature for about 3 hours. After that, the mixture was centrifuged at 5000 rpm for 5 min twice, and the excess of transferrin was discarded. The pellet was resuspended in 100 µL of 2 mM borate buffer (pH 8.5).

### Cell culture for live cell imaging

A 549 human lung cancer cells (CCL-185, ATCC) were cultured on a clean 22× 22 mm coverslip in a plastic petri dish. The cells were maintained in complete cell culture containing Dulbecco’s Modified Eagle’s Medium (DMEM) and 10% fetal bovine serum (FBS, 30-2020, ATCC) at 37 °C, 5% CO_2_ in a humidified atmosphere. For the live cell imaging experiment, two pieces of double-sided tape were attached on top of the clean glass slide. A coverslip with adherent cells was put on top of the taps to form a chamber, with the cell side facing the glass slide. 8 µL transferrin@GNRs mixed with 42 µL cell culture without FBS were injected into the chamber, and the chamber was sealed by nail polish.

### Microtubule labeling

To monitor the microtubule in live cell, the microtubules were labeled by SiR700-tubulin Kit (Cytoskeleton) at a final concentration of 800 nM in cell culture medium for 2 h before imaging.

### Optical imaging

All optical imaging experiments were performed on a Nikon 80i microscope equipped with a 100W halogen tungsten lamp. When the microscopy was set in Dark-filed mode, A 650 nm band-pass filter was placed between the light source and sample. The parallax dark field mode was achieved by inserting a half wedge prism (0.5° Edmund, Barrington, NJ) to the light path. For dark field microscopy, a numerical aperture 1.20-1.43 oil immersion dark field condenser, and a 60× Plan fluor oil immersion objective was used. An additional 2× magnification was used to result in a pixel size of 131 nm. An extra lens with focusing length of 500 mm was inserted in the defocused channel before the detector so that the axial separation distance between the focal planes in two channels was fixed to 0.9 µm. When the microscopy was switched to epifluorescence mode, microtubule labeled by Sir700-tubulin kit was imaged with standard Cy5 filter set. 100-200 images from the same sample were obtained and then applied the SRRF analysis.

Images and movies were recorded by an EMCCD camera (Andor iXon Ultra 897). The focal plane was locked to the target probe by coupling with an objective scanner (PI, PD72Z1) to adjust the z-position of the sample. The correlation mapping and calibration algorithm were compiled in a plug-in for ImageJ µManager to operate the imaging system. All scattering images were recorded at 50 frames per second (fps), while fluorescence images were monitored before or after one intracellular transport event with exposure time of 0.5 s. A heating stage was used to maintain the physiological temperature of 37°C. The acquired movies were analyzed using ImageJ and MATLAB.

## Results

### Bifocal parallax SPT

Bifocal imaging (37), parallax microscopy (38-40), and an automatic feedback control module are the key components of the imaging system (**Fig. 1A**) for 3D defocused SPT. The design principle is to keep the target probe in focus continuously in one channel to provide high sensitivity for extracting the 3D spatial coordinates (x, y, z) with nanometer precision, while *maintaining a constant defocusing distance* in another channel for determining the probe’s orientation angles accurately and robustly from image pattern matching.

**Figure 1.**
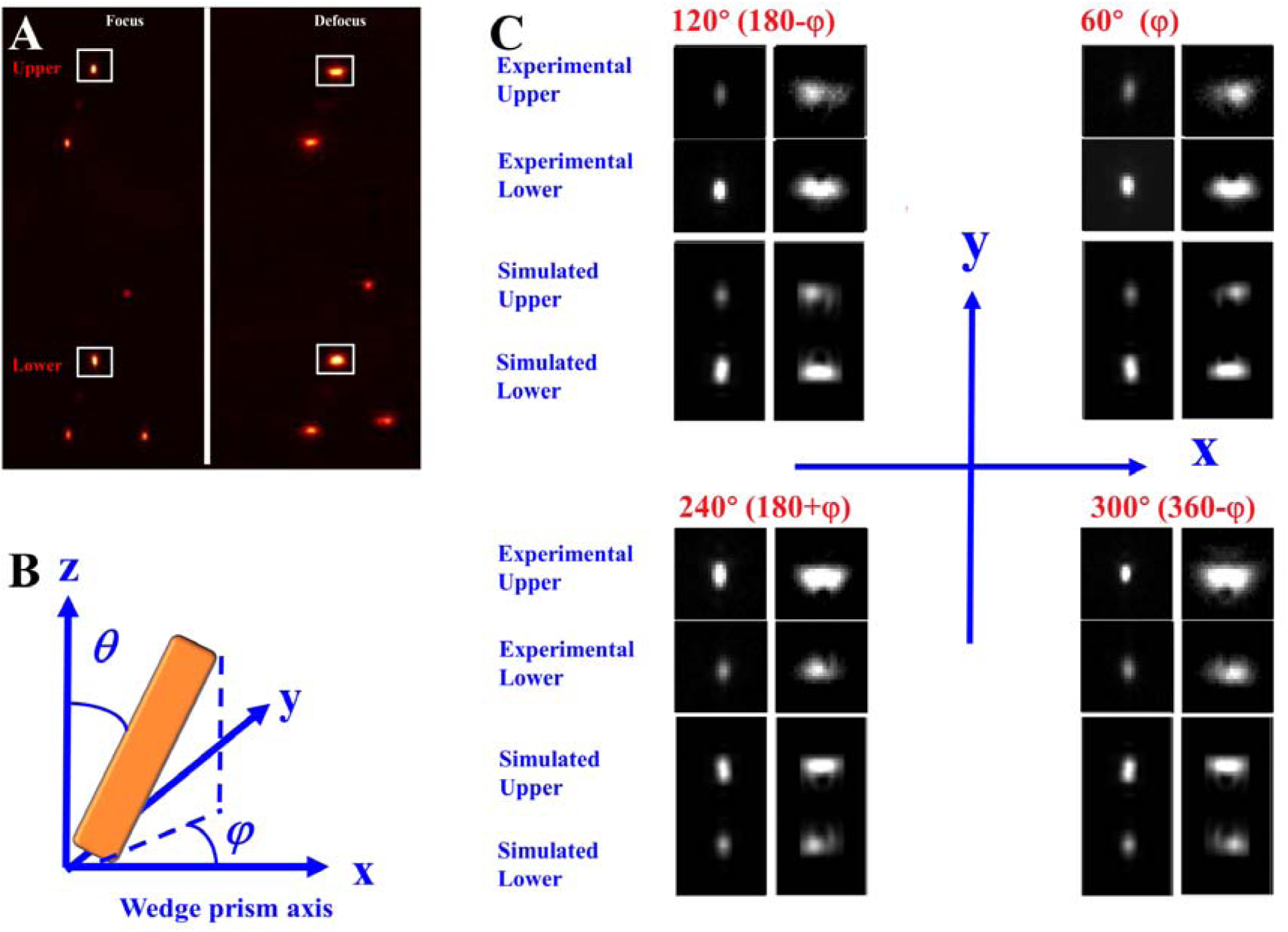
Bifocal parallax SPT. (A) Sample of experimental image. The white line squares label a group of four spots (upper and lower in-focus and defocused patterns). The upper and lower in-focus spots have the same x coordinate and the distance. The defocusing distance for the defocused channel is 0.9 μm. Scale bar is 10 μm. (B) Definitions of azimuth angle *φ* and polar angle *θ* of a GNR with respect to wedge prism axis. (C) Experimental and simulated half-plane scattering images (upper and lower in-focus spots, upper and lower defocus patterns) of a GNR with polar angle of 60° as a function of azimuth angle every 60°. Scale bar is 2 μm.

In bifocal imaging, a beam splitter (reflection:transmission = 70:30) was used to split the collected signal into two light paths and create two imaging channels. An extra lens with a focusing length of 500 mm was inserted in one channel (70%) to generate defocused imaging pattern. An axial separation distance of 0.9 μm which was found to generate the most suitable defocused image patterns (22) (**Fig. S1**) between two imaging channels (focused vs. defocused) was introduced and locked during the imaging experiments.

For parallax imaging, a wedge prism was inserted at the back focal plane of the objective to deviate half of the scattering light from the sample by a very small angle, effectively splitting the light to form two *half-plane* images (as opposed to the standard full-plane images using the full pupil) in both focused and defocused channels. The two associated half-plane mirror images for a single probe are always aligned vertically, and they will be referred to as the *upper* and *lower* image, respectively, in the rest of discussions. A calibration curve was constructed experimentally to correlate the distance between the two mirror spots in the x-y plane (Δy) with the vertical position (z) of the target probe (***SI Text*** and **Fig. S2**).

Finally, the automatic feedback control system, composed of a piezo objective scanner and a tracking program in µManger (41, 42), was implemented to keep the target probe in focus. When the target probe was in focus, the y distance between the upper and lower images in the focused channel was obtained as the reference (Δy_0_). During the tracking, Δy was calculated for every frame. Any axial movement of the target probe would give changes to the measured Δy, and the tracking program would respond by moving the objective scanner accordingly to keep the target probe in focus.

This bifocal parallax SPT system generates a total of four images (**Fig. 1A**, the focused upper and lower images on the left and the defocused upper and lower images on the right) of the target probe on the same electron multiplying charged-coupled device (EMCCD) camera to provide all the information we need to resolve its detailed translational and rotational motions. The x and y coordinates can be resolved by correlation mapping in the focused channel, and the z coordinate can be obtained from the recorded movement of the objective scanner. The experimental localization precisions in the x-, y- and z-axes were measured to be 2.2 ± 0.1, 4.8 ± 0.3, and 11.6 ± 0.5 nm, respectively (***SI Text*** and **Fig. S3**). The localization precision is worse in y compared to that in x because the half-plane point spread functions (PSFs) are stretched in y in our imaging setup.

### Determining 3D orientation

The orientation angles are defined in **Fig. 1B**. Examples of the half-plane scattering images of a 40 nm × 80 nm GNR immobilized in agarose gel with a fixed polar angle of 60° and various azimuth angles are shown in **Fig. 1C**, and a complete set of images with an azimuth angle interval of 10° is provided as **Fig. S4**.

Computer simulation of PSF is essential in acquiring a complete understanding of the experimental image patterns and to estimate uncertainties in the related measurements. Several methods have been developed to determine the dipole orientation by fitting experimental images with the simulated patterns (43, 44). However, none of the existing simulation programs can be used directly for parallax microscopy, which employs a semi-circle pupil to result in the half-plane PSFs. To fully understand the irregular shapes of the defocused half-plane images, a simulation program to obtain theoretical half-plane PSFs as a function of azimuth and polar angles using the six basic functions of dipole emission 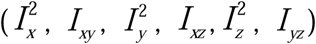 (45, 46) has been developed in house (see the SI text for details on simulation). By introducing two semi-circle pupils for the upper and lower half-plane PSFs, respectively, the six basic patterns are transformed into two groups of basic half-plane patterns (**Fig. S5**). These basic patterns are dependent on system-specific parameters, including the numerical aperture, magnification of the objective, and defocusing distance, and their linear combination can reproduce the half-plane image patterns.

The simulated upper and lower half-plane defocused images of four different azimuth angles (φ, 180°-φ, 180°+φ, and 360°-φ) and a same polar angle agree well with the experimental counterparts (**Fig. 1C**). More importantly, the defocused image patterns are clearly unique at these azimuth angles in the four quadrants of the Cartesian plane. Therefore, the half-plane PSFs in defocused parallax microscopy, like the conventional (full-plane) defocused PSFs, can determine azimuth angles without angular degeneracy.

A complete set of simulated images with various combinations of azimuth and polar angles was used to build a database for pattern matching, an established strategy for determining the 3D orientation from defocused images (19). The analysis of the orientation-dependent uncertainties associated with the estimated azimuth and polar angles is provided in **Fig. S6** and **S7**. Under our typical live cell imaging conditions, a precision of < 2° can be accomplished for most polar and azimuth angles with a signal to noise ratio of 10.

### Correlation of transport trajectories with microtubule geometry

To use GNRs as reporter of the rotational motion in living cells, GNRs must be first introduced into the cells. It has been demonstrated previously that transferrin-modified GNRs were internalized mainly through the clathrin-mediated endocytosis pathway in cells expressing transferrin receptors in the cell membrane (including A549 human lung cancer cells used in the current study) (47, 48). In these GNR-containing early endosomes, the GNR is wrapped tightly by the membrane and show no significant movement relative to the vesicle, which meets the criterion for rotational tracking (16).

Once inside a cell, the cargo can be transported by molecular motors on the cytoskeleton. The 3D organization, especially the intersections, of the microtubules can dramatically affect the transport behaviors. The axial separation distance and the angle between intersecting cytoskeleton filaments can maintain, block or change a cargo’s moving direction (34, 49, 50). In order to establish a correlation between the 3D organization of microtubule tracks and the dynamics of transporting cargoes, we labeled the microtubule network in living A549 cells with the SiR700-tubulin kit, which consists of silicon rhodamine fluorophore and the microtubule-binding drug docetaxel and has been proved to have little impact on cell proliferation even at concentrations up to 100 nM(51) and was widely used to study the motor protein motility (52-54). We then applied the Super-Resolution Radial Fluctuations (SRRF) analysis, which is capable of revealing high-fidelity super-resolution information with ∼ 150 nm resolution in conventional epifluorescence microscopy images (55). In coSmbination with the 3D trajectories of cargoes, the detailed 3D geometry of microtubule can be uncovered (**Fig. S8**). Compared to the previously reported approach of super-resolution imaging after cell fixation (30), SRRF avoids potential problems of introducing dynamic structural changes of microtubule network during cell fixation. On the other hand, SRRF offers sufficient spatial resolution to distinguish the individual microtubules involved in the recorded cargo transport events.

### Characterizing 3D free diffusion and active transport

Two basic phases of the motor-driven intracellular transport exist in every recorded trajectory: the active transport phase in which the cargo maintains a relatively constant transport velocity, and the pause phase in which the cargo has little or no translational movement (18). Pauses frequently happen at the microtubule intersections (30, 50), as well as at damaged microtubules (56) and microtubule-actin (57) or actin-actin (58) junctions. These observations are consistent with translation-based SPT.

When the rotational dimensions are included in the observation, the pause phase can be further classified into two groups, those with minimal rotation and with active rotation. Notably, they are not distinguishable in translation-based SPT. To better understand the cargo’s motion at the pause phase and its implication on the cargo-motor-track interaction dynamics in living cells, in this section we will first discuss the well-understood cases of *active transport* and *free diffusion*, which sets the foundation for further studies of the more complex cases that are closely related to how cargoes move across different microtubule geometries. Examples of free diffusion were obtained in cells treated with colchicine to induce microtubule disassembly (59).

Key parameters for different movement patterns are compared in **Fig. 2** and **Table S1**. Active transport and free diffusion can be readily distinguished by the measured exponent α in mean squared displacement (MSD) analysis (60). As expected, the active transport examples in our SPT experiments gave a characteristic exponent α much great than 1, suggesting a super-diffusion model; on the other hand, the measured exponent α for free diffusion is close to 1. It is worthwhile to note that unlike most previous studies in which 2D diffusion was analyzed in the horizontal x-y plane, our imaging method allows the recording of movement along the z-axis and therefore the measurement of 3D diffusion in living cells. **Fig. S9** shows an active transport example of a GNR-containing cargo moving along the fluorescently-labeled microtubules with a vertical movement of ∼300 nm.

**Fig. 2.**
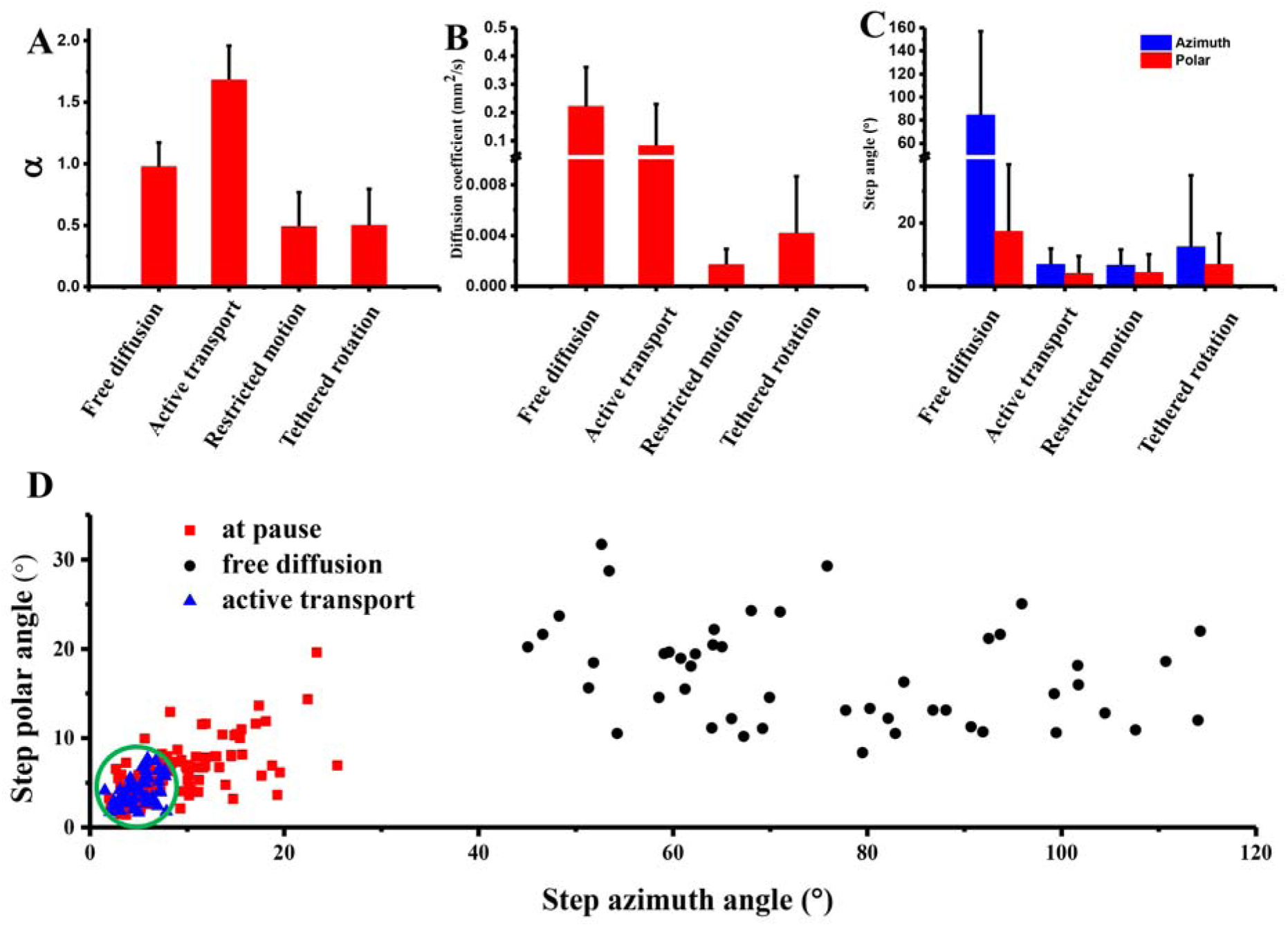
Parameters for describing the motion of GNR cargoes at free diffusion, active transport, tight attachment and tethered rotation. (A) The exponent α of mean squared displacement. Each data point is >50 recorded events. (B) Apparent diffusion coefficients (>50 events each). (C) Step azimuth angle and step polar angle (>1500 steps each). (D) The scatter plot of the step polar and azimuth angles calculated for individual recorded trajectories. The green circle indicates the boundary of data points for the cases of active transport and tight attachment. Its radius is 2.5x the standard deviation of the active transport cases (blue triangles).

The exponent α value is commonly used in the analysis of diffusion; however, it alone is insufficient to provide the detailed dynamics information. The recorded rotational motion can be described quantitatively using step azimuth angle (Δφ) and step polar angle (Δθ), which are defined as the angular difference in two consecutive frames (20 ms time interval). For active transport, the average step azimuth and polar angles were small (< 5°, **Fig. 2C**), suggesting the cargoes were attached firmly to the microtubule by kinesin and dynein motors with greatly restrained angle ranges. The detected subtle angle changes can be attributed to thermal fluctuations and the shake of stalk caused by motor stepping. For free diffusion, the average step azimuth and polar angles were significantly larger, indicating more freedom for the cargo to rotate in broad angle ranges.

### Classification of cargo-motor-track interaction at pauses

Next, the parameters are used to distinguish more complicated cases at pauses. **Fig. 2D** is a scatter plot of the step azimuth and polar angles for the pauses (red squares) with each data point represents one recorded event. As a comparison, the same parameters for active transport (blue triangles) and free diffusion (black squares) are also plotted. Visually, with the 2.5x the standard deviation of step angle of the active transport cases as boundary, two modes of cargo-motor-track interaction at pauses can be identified from this distribution.

The first mode is named *tight attachment*, during which the cargo can only rotate in narrow angle ranges, likely because the cargo, motor and microtubule are fully engaged with the tensions applied by multiple motor proteins from multiple directions to restrict the lateral movement and rotational motion of the cargo. The step azimuth and polar angles for the tight attachment cases are similar to those during active transport (inside the green circle in **Fig. 2D**).

The second mode is named *tethered rotation*, during which the cargo is also attached to the microtubule by showing a lack of translational freedom; however, the binding between the cargo and the microtubule is loose, therefore giving more rotational freedom to the cargo. The step azimuth and polar angles for the tethered rotation cases (red squares outside of the green circle in **Fig. 2D**) are significantly smaller than those of free diffusion but larger than those of active transport and tight attachment. It is conjectured that the cargo is attached to the microtubule from one direction, which lead to the chain hammer-like movement of the cargo. Furthermore, the exponent α of ∼0.5 for both *tight attachment* and *tethered rotation* implies a sub-diffusion model, suggesting restricted translational motions at the pauses.

Clearly, by quantitatively analyzing the step azimuth and polar angle, we can describe the interactions between the cargo, motor and track more precisely and in greater detail than all previous methods. Based on the understanding of the basic motion modes at pauses, in the following sections, we will elucidate the cargo transport dynamics demonstrating these motions at the microtubule intersections.

### Representative dynamic behaviors of cargo at microtubule intersections

The capability of revealing the full spatial information (including the 3D spatial coordinates and orientation angles) of the GNR-containing cargo and the dynamic cargo-motor-track interaction is essential in visualizing how the cargo adapts its posture to the underlying track geometry through the interactions with the motors and microtubules and gaining new insights on the mechanism of intracellular transportation on the complex 3D microtubule network. *Two types of distinct dynamic behaviors* of the cargo at the microtubule intersections have been identified.

#### Type I: Tight attachment througout followed by active transport

This type of dynamic behavior is characterized with tight attachment at the pauses at the interactions. It is conjectured that the cargo is bound to the microtubule, or possibly multiple surrounding microtubules, through the kinesin and dynein motors. Individual motors may detach from and reattach to the microtubule(s), allowing the cargo to adjust its position slightly (∼ a few nm), either following the original or a new track; however, tensions exist due to the competing, engaged motors from multiple directions that restrain the cargo’s rotation. Then, when either kinesin or dynein motors win the “tug-of-war”, the winning motors exert forces on the cargo with a pulling motion. The cargo resumes to the active transport phase, often accompanied by a sudden change of orientation.

**Fig. 3** shows such an example that the cargo experienced an apparent pause phase, during which it actually finished a U-turn on a microtubule intersection consisting of three microtubules revealed in the SRRF image: two nearly parallel microtubules *a, c* with a separation distance of ∼ 130 nm and another intersecting microtubule *b*. The three microtubules were in close proximity at the intersection to allow the simultaneous attachment of the cargo by the motors and a smooth, nearly nonstop transition from microtubule *a* to microtubule *c*. The recording and reconstitution of this event are provided as **Movies S1 and S2**, respectively.

**Fig. 3.**
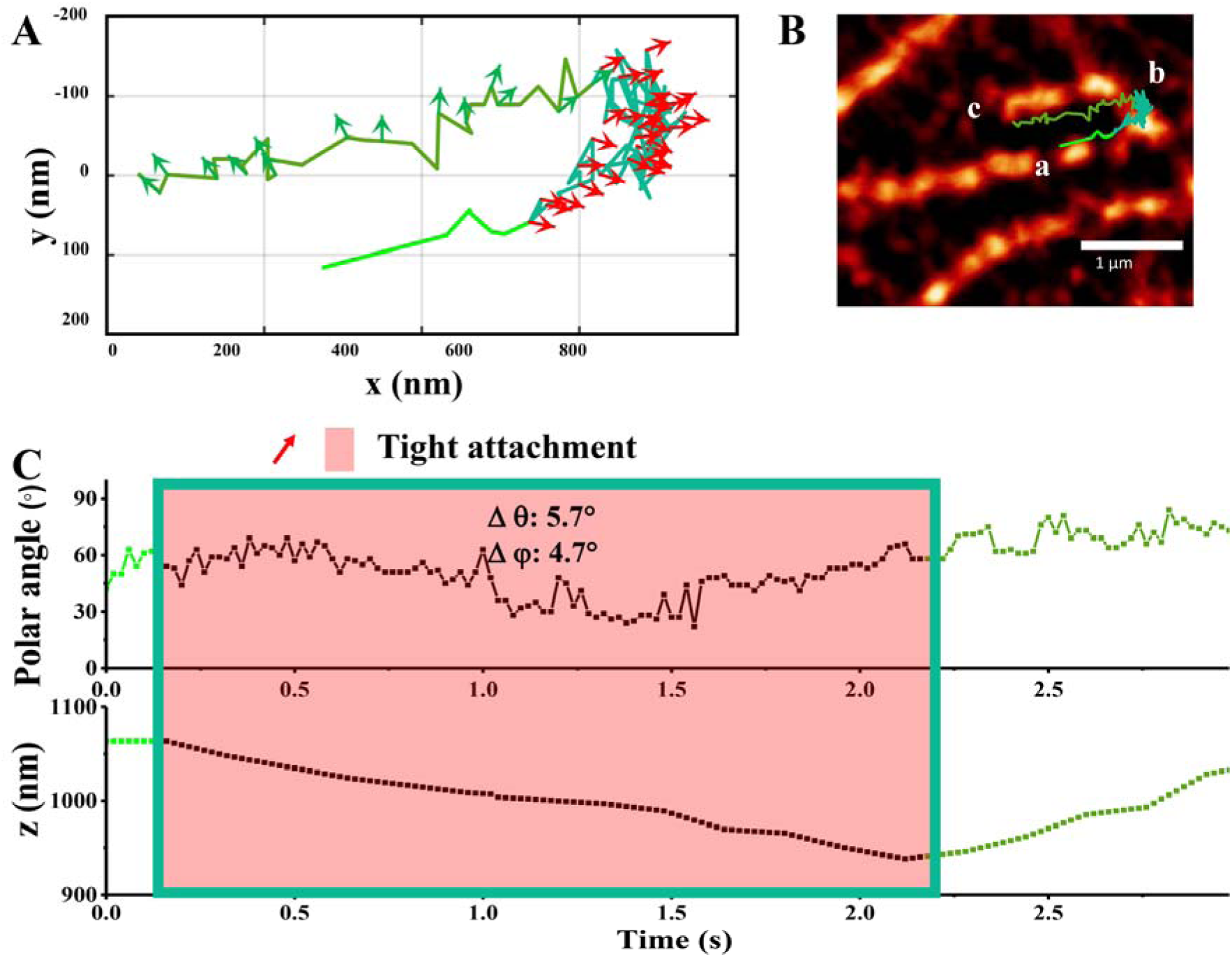
Translational and rotational motion of a cargo crossing a U-shape microtubule intersection. (A) 2D trajectory. The azimuth angle in every three frames (a time interval of 60 ms) is displayed as an arrow (red: tight attachment: gray: a quick release from tension (azimuth angle changing by ∼90° counterclockwise). (B) Overlap of the cargo’s trajectory and the SRRF image of the microtubules: two nearly parallel microtubules *a, c* with a separation distance of ∼ 130 nm and another intersecting microtubule *b*. (C) Time series of the polar angle and z displacement. The inserted Δφ, Δθ are the average step azimuth angle and polar angle for the entire red zone. The solid line in the top panel guides the changes of the polar angle over time.

In the early stage of this process, the cargo took active transport until reaching the intersection of the three microtubules. Then the cargo paused while undergoing a tight attachment (the red zone in **Fig. 3**) with small azimuth angle changes (average Δφ = 5.8°). The polar angle changed gradually (from ∼60° to ∼25° and then back to ∼60°) while moving downward ∼130 nm along microtubule *b*, likely being pulled by the motors attached to microtubule *b* while adapting its posture to suit the track geometry. In the final moment of the pause, the cargo experienced a quick release from tension (azimuth angle changing by ∼90° counterclockwise) while moving upward ∼100 nm along microtubule *c*.

#### Type II: Switching between tight attachment and tethered rotation repeatedly

The second type of dynamic behavior is defined by *alternated tight attachment and tethered rotation*. Similar to Type I, when the cargo is in the *tight attachment* mode, it is also possibly bound to the microtubule(s) by different types of motor proteins. However, it appears that micro-tuning the position of the cargo is insufficient for the cargo to go over (or under) the obstacle or to find the new track. Thus, there is the *tethered rotation* mode, which is characterized by significantly larger angle ranges (in other words, more rotational freedom). This binding status of the cargo can be reflected in its relatively large step azimuth and polar angles, as well as large z fluctuations (∼100 nm). *Tethered rotation* gives the cargo more flexibility to search for and connect to new microtubules, while *tight attachment* corresponds to the status that the cargo has found and connected to the surrounding microtubules and the motors have started to engage. The steric hindrance and the pulling force from the lagging motors may contribute to the detachment of the leading motors, which allows them to search for a new possible path.

An example of Type II is shown in **Fig. 4** and **Movie S3**. During the first part of the long pause (0.68 – 6.08 s) of this example, the cargo underwent alternate tight attachment and tethered rotation with only small oscillating z movement, suggesting the transient binding of the motors to the surrounding microtubules. Then, the cargo went into an extended tethered rotation state (6.08-8.34 s) with larger step azimuth and polar angle changes while moving vertically for a distance of ∼100 nm, suggesting a search for microtubule binding in a wider region. In the final period (8.34-10.54 s), the cargo experienced another tight attachment with gradual azimuth angle change (a ∼150° counterclockwise rotation followed by another ∼150° clockwise rotation; see **Fig. S10** for experimental and matching simulated image patterns for this directional rotation), likely a process for kinesin motors to win the competition and drive the cargo onto the crossing track toward the nucleus.

**Fig. 4.**
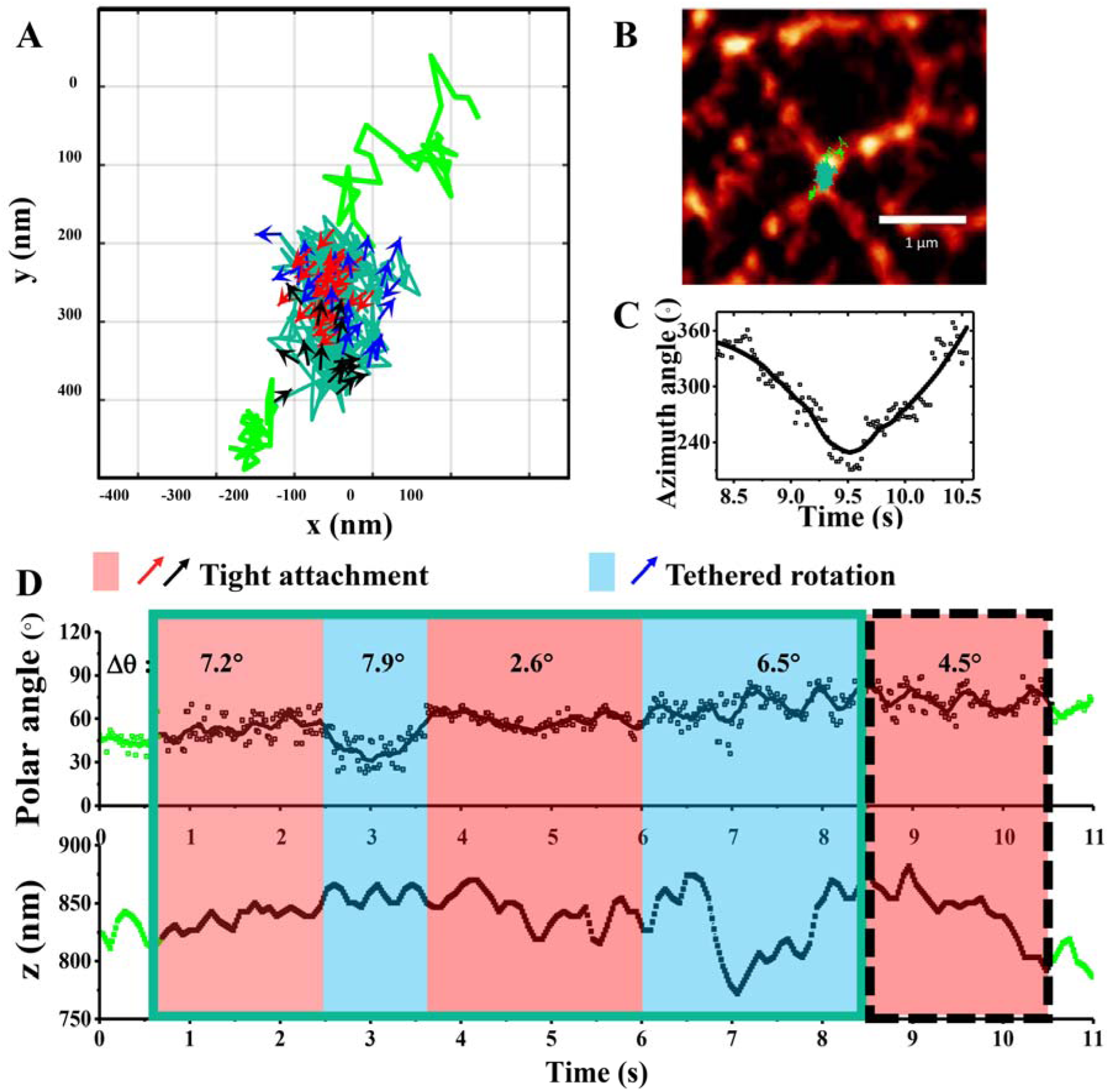
Translational and rotational motion of a cargo switching between tight attachment and tethered rotation repeatedly at pause. (A) 2D trajectory with the color-coded arrows (every ten frames or 200 ms) indicating the azimuth angle of the GNR (blue for tethered rotation; red/black for tight attachment). The black arrows demonstrate a special case of tight attachment during which a directed rotation was observed. (B) Overlap of the 2D trajectory and the SRRF image of microtubules. (C) The azimuth angle of GNR as a function of time for the directed rotation (8.34-10.54 s). (D) Polar angle and z of the GNR as a function of time. The inserted Δφ, Δθ indicate step azimuth angle and polar angle of various stages. The red and green shadow indicates the tight attachment and tethered rotation periods, respectively, and the black dashed square indicates the final period of the directed rotation.

Alternating between tight attachment and tethered rotation was commonly found at the pauses. As shown in **Fig. S11**, ∼90% of the recorded pauses experienced more than 1 switch. As expected, there is also a positive correlation between the number of switching and the pause duration. The cargoes often need longer time and rearrange their motor several times to adjust their posture to suit for the microtubule geometry to overcome the pause. This type of behavior possibly represents a practically more challenging case that requires the cargo to partially detach from the microtubule to get over the obstacle or find its new path.

## Discussion and outlook

The bifocal parallax SPT method presented here harvests the full potential of defocused imaging for dynamic 3D tracking of anisotropic imaging probes in complex cellular environments by having a reliable way to maintain a fixed defocusing distance and therefore accurate measurement of dipole orientation, while keeping track of any movement in the z-axis.

Using this method, we have resolved the highly detailed translational and rotational dynamics of GNR-containing cargo transporting at the microtubule intersections in living cell. The quantitatively resolved rotational motion of the cargo is demonstrated to be highly insightful in revealing the dynamics of the cargo-motor-track interactions.

The two basic rotation modes of cargoes at pauses may be indicative of the status of motor-cargo-microtubule interactions. Tight attachment is likely associated with the competition of motors, in which motors are engaged to apply tensions on the cargo from multiple directions and restrict the cargo’s rotation. On the other hand, tethered rotation represents a more flexible state of the cargo searching for the next transport opportunity through coordinated behavior of motors, possibly through stress-associated detachment. Cargo position adjustment and motor binding to/unbinding from the microtubules are involvement in both modes. Correlating cargo dynamics with coordination factors may be able to provide important mechanistic insights on how the cargo adapts to the local environment through modulation of the interactions among the cargo, motor, and microtubule in future studies.

The current study also demonstrates the potential of studying coordinated interactions in living cellular environments from resolving characteristic rotational motions. Surface-modified gold nanoparticles can be employed as a model system to interrogate nanoparticle-cell interactions. This method is also generally applicable to the studies of membrane processes, including adhesion (61) (sample question: how do objects of different shapes approach the membrane?), diffusion (62) (how does a target make transitions back and forth between confined diffusion and hopping diffusion?), membrane protein binding (63) (what is the correlation of the motions of nanoprobes with their binding status to membrane receptors?), endocytosis (64) (how does the dynamin helices cut vesicles at the end of clathrin-mediated endocytosis?), virus infection (65) (how does the virus enter the cell and control the motor protein to transport them to their destinatonin to infect the cell?), etc.

## Author Contributions

X.C., K.C. and N.F. designed research. X.C., K.C. and B.D. built the imaging setup and performed the imaging experiments. X.C. and S.L.F. prepared the nanoparticle samples. All authors wrote the manuscript.

## Acknowledgments

The instrument and method development were supported by the National Science Foundation (CBET-1604612). The biophysical study of cellular processes was supported by the National Institution of Health (R01GM115763).

## Competing Interests

The authors declare no competing interests.

## Notes

### Competing Interest Statement

The authors have declared no competing interest.

## References

1. Pollard, T. D. 2010. A guide to simple and informative binding assays. Mol. Biol. Cell 21(23):4061–4067.

2. Rasmussen, S. G., H.-J. Choi, D. M. Rosenbaum, T. S. Kobilka, F. S. Thian, P. C. Edwards, M. Burghammer, V. R. Ratnala, R. Sanishvili, and R. F. Fischetti. 2007. Crystal structure of the human β 2 adrenergic G-protein-coupled receptor. Nature 450(7168):383.

3. Stark, H., and R. Lührmann. 2006. Cryo-electron microscopy of spliceosomal components. Annu. Rev. Biophys. Biomol. Struct. 35:435–457.

4. Neuman, K. C., and A. Nagy. 2008. Single-molecule force spectroscopy: optical tweezers, magnetic tweezers and atomic force microscopy. Nat. Methods 5(6):491.

5. Manley, S., J. M. Gillette, G. H. Patterson, H. Shroff, H. F. Hess, E. Betzig, and J. Lippincott-Schwartz. 2008. High-density mapping of single-molecule trajectories with photoactivated localization microscopy. Nat. Methods 5(2):155.

6. Jaqaman, K., D. Loerke, M. Mettlen, H. Kuwata, S. Grinstein, S. L. Schmid, and G. Danuser. 2008. Robust single-particle tracking in live-cell time-lapse sequences. Nat. Methods 5(8):695.

7. Liang, L., J. Li, Q. Li, Q. Huang, J. Shi, H. Yan, and C. Fan. 2014. Single-particle tracking and modulation of cell entry pathways of a tetrahedral DNA nanostructure in live cells. Angew. Chem. Int. Edit. 53(30):7745–7750.

8. Sanamrad, A., F. Persson, E. G. Lundius, D. Fange, A. H. Gynnå, and J. Elf. 2014. Single-particle tracking reveals that free ribosomal subunits are not excluded from the Escherichia coli nucleoid. Proc. Natl Acad. Sci. USA 111(31):11413–11418.

9. von Diezmann, A., Y. Shechtman, and W. Moerner. 2017. Three-dimensional localization of single molecules for super-resolution imaging and single-particle tracking. Chem. Rev. 117(11):7244–7275.

10. Adachi, K., K. Oiwa, T. Nishizaka, S. Furuike, H. Noji, H. Itoh, M. Yoshida, and K. Kinosita Jr. 2007. Coupling of rotation and catalysis in F1-ATPase revealed by single-molecule imaging and manipulation. Cell 130(2):309–321.

11. Roux, A., K. Uyhazi, A. Frost, and P. De Camilli. 2006. GTP-dependent twisting of dynamin implicates constriction and tension in membrane fission. Nature 441(7092):528.

12. Backlund, M. P., A. Arbabi, P. N. Petrov, E. Arbabi, S. Saurabh, A. Faraon, and W. Moerner. 2016. Removing orientation-induced localization biases in single-molecule microscopy using a broadband metasurface mask. Nat. Photonics. 10(7):459.

13. Forkey, J. N., M. E. Quinlan, M. A. Shaw, J. E. Corrie, and Y. E. Goldman. 2003. Three-dimensional structural dynamics of myosin V by single-molecule fluorescence polarization. Nature 422(6930):399.

14. Sönnichsen, C., and A. P. Alivisatos. 2005. Gold nanorods as novel nonbleaching plasmon-based orientation sensors for polarized single-particle microscopy. Nano Lett. 5(2):301–304.

15. Xu, D., Y. He, and E. S. Yeung. 2014. Direct Observation of the Orientation Dynamics of Single Protein-Coated Nanoparticles at Liquid/Solid Interfaces. Angew. Chem. Int. Edit. 53(27):6951–6955.

16. Kaplan, L., A. Ierokomos, P. Chowdary, Z. Bryant, and B. Cui. 2018. Rotation of endosomes demonstrates coordination of molecular motors during axonal transport. Sci. Adv 4(3):e1602170.

17. Wang, G., W. Sun, Y. Luo, and N. Fang. 2010. Resolving rotational motions of nano-objects in engineered environments and live cells with gold nanorods and differential interference contrast microscopy. J. Am. Chem. Soc. 132(46):16417–16422.

18. Gu, Y., W. Sun, G. Wang, K. Jeftinija, S. Jeftinija, and N. Fang. 2012. Rotational dynamics of cargos at pauses during axonal transport. Nat. Commun. 3:1030.

19. Toprak, E., J. Enderlein, S. Syed, S. A. McKinney, R. G. Petschek, T. Ha, Y. E. Goldman, and P. R. Selvin. 2006. Defocused orientation and position imaging (DOPI) of myosin V. Proc. Natl Acad. Sci. USA 103(17):6495–6499.

20. Lew, M. D., and W. Moerner. 2014. Azimuthal polarization filtering for accurate, precise, and robust single-molecule localization microscopy. Nano Lett. 14(11):6407–6413.

21. Li, T., Q. Li, Y. Xu, X.-J. Chen, Q.-F. Dai, H. Liu, S. Lan, S. Tie, and L.-J. Wu. 2012. Three-dimensional orientation sensors by defocused imaging of gold nanorods through an ordinary wide-field microscope. ACS Nano 6(2):1268–1277.

22. Xiao, L., Y. Qiao, Y. He, and E. S. Yeung. 2010. Three dimensional orientational imaging of nanoparticles with darkfield microscopy. Anal. Chem. 82(12):5268–5274.

23. Schuster, M., R. Lipowsky, M.-A. Assmann, P. Lenz, and G. Steinberg. 2011. Transient binding of dynein controls bidirectional long-range motility of early endosomes. Proc. Natl Acad. Sci. USA 108(9):3618–3623.

24. Ananthanarayanan, V., M. Schattat, S. K. Vogel, A. Krull, N. Pavin, and I. M. Tolić-Nørrelykke. 2013. Dynein motion switches from diffusive to directed upon cortical anchoring. Cell 153(7):1526–1536.

25. Hancock, W. O., and J. Howard. 1998. Processivity of the motor protein kinesin requires two heads. J. Cell Biol. 140(6):1395–1405.

26. Schlager, M. A., and C. C. Hoogenraad. 2009. Basic mechanisms for recognition and transport of synaptic cargos. Mol. Brain 2(1):25.

27. Zajac, A. L., Y. E. Goldman, E. L. Holzbaur, and E. M. Ostap. 2013. Local cytoskeletal and organelle interactions impact molecular-motor-driven early endosomal trafficking. Curr. Biol. 23(13):1173–1180.

28. Müller, M. J., S. Klumpp, and R. Lipowsky. 2008. Tug-of-war as a cooperative mechanism for bidirectional cargo transport by molecular motors. Proc. Natl Acad. Sci. USA 105(12):4609–4614.

29. Belyy, V., M. A. Schlager, H. Foster, A. E. Reimer, A. P. Carter, and A. Yildiz. 2016. The mammalian dynein–dynactin complex is a strong opponent to kinesin in a tug-of-war competition. Nat. Cell Biol. 18(9):1018.

30. Osunbayo, O., J. Butterfield, J. Bergman, L. Mershon, V. Rodionov, and M. Vershinin. 2015. Cargo transport at microtubule crossings: evidence for prolonged tug-of-war between kinesin motors. Biophys. J. 108(6):1480–1483.

31. Hancock, W. O. 2014. Bidirectional cargo transport: moving beyond tug of war. Nat. Rev. Mol. Cell Bio 15(9):615–628.

32. Ross, J. L., H. Shuman, E. L. Holzbaur, and Y. E. Goldman. 2008. Kinesin and dynein-dynactin at intersecting microtubules: motor density affects dynein function. Biophys. J. 94(8):3115–3125.

33. Schroeder III, H. W., A. G. Hendricks, K. Ikeda, H. Shuman, V. Rodionov, M. Ikebe, Y. E. Goldman, and E. L. Holzbaur. 2012. Force-dependent detachment of kinesin-2 biases track switching at cytoskeletal filament intersections. Biophys. J. 103(1):48–58.

34. Bergman, J. P., M. J. Bovyn, F. F. Doval, A. Sharma, M. V. Gudheti, S. P. Gross, J. F. Allard, and M. D. Vershinin. 2018. Cargo navigation across 3D microtubule intersections. Proc. Natl Acad. Sci. USA:201707936.

35. Verdeny-Vilanova, I., F. Wehnekamp, N. Mohan, Á. S. Álvarez, J. S. Borbely, J. J. Otterstrom, D. C. Lamb, and M. Lakadamyali. 2017. 3D motion of vesicles along microtubules helps them to circumvent obstacles in cells. J. Cell Sci. 130(11):1904–1916.

36. Filbrun, S. L., A. B. Filbrun, F. L. Lovato, S. H. Oh, E. A. Driskell, and J. D. Driskell. 2017. Chemical modification of antibodies enables the formation of stable antibody–gold nanoparticle conjugates for biosensing. Analyst. 142(23):4456–4467.

37. Toprak, E., H. Balci, B. H. Blehm, and P. R. Selvin. 2007. Three-dimensional particle tracking via bifocal imaging. Nano Lett. 7(7):2043–2045.

38. Chen, K., Y. Gu, W. Sun, G. Wang, X. Fan, T. Xia, and N. Fang. 2017. Characteristic rotational behaviors of rod-shaped cargo revealed by automated five-dimensional single particle tracking. Nat. Commun. 8(1):887.

39. Sun, Y., J. D. McKenna, J. M. Murray, E. M. Ostap, and Y. E. Goldman. 2009. Parallax: high accuracy three-dimensional single molecule tracking using split images. Nano Lett. 9(7):2676–2682.

40. Yajima, J., K. Mizutani, and T. Nishizaka. 2008. A torque component present in mitotic kinesin Eg5 revealed by three-dimensional tracking. Nat. Struct. Mol. Biol 15(10):1119.

41. Schneider, C. A., W. S. Rasband, and K. W. Eliceiri. 2012. NIH Image to ImageJ: 25 years of image analysis. Nat. Methods 9(7):671.

42. Edelstein, A., N. Amodaj, K. Hoover, R. Vale, and N. Stuurman. 2010. Computer control of microscopes using µManager. Curr. Protoc. Mol. Biol. 92(1):14.20. 11-14.20. 17.

43. Patra, D., I. Gregor, and J. Enderlein. 2004. Image analysis of defocused single-molecule images for three-dimensional molecule orientation studies. J. Phys. Chem. A 108(33):6836–6841.

44. Aguet, F., S. Geissbühler, I. Märki, T. Lasser, and M. Unser. 2009. Super-resolution orientation estimation and localization of fluorescent dipoles using 3-D steerable filters. Opt. Express 17(8):6829–6848.

45. Burghardt, T. P. 2011. Single molecule fluorescence image patterns linked to dipole orientation and axial position: application to myosin cross-bridges in muscle fibers. PLoS One 6(2):e16772.

46. Backer, A. S., and W. Moerner. 2014. Extending single-molecule microscopy using optical Fourier processing. J. Phys. Chem. B 118(28):8313–8329.

47. McMahon, H. T., and E. Boucrot. 2011. Molecular mechanism and physiological functions of clathrin-mediated endocytosis. Nat. Rev. Mol. Cell Bio 12(8):517.

48. Chithrani, B. D., and W. C. Chan. 2007. Elucidating the mechanism of cellular uptake and removal of protein-coated gold nanoparticles of different sizes and shapes. Nano Lett. 7(6):1542–1550.

49. Ross, J. L., M. Y. Ali, and D. M. Warshaw. 2008. Cargo transport: molecular motors navigate a complex cytoskeleton. Curr. Opin. Cell Biol. 20(1):41–47.

50. Bálint, Š., I. V. Vilanova, Á. S. Álvarez, and M. Lakadamyali. 2013. Correlative live-cell and superresolution microscopy reveals cargo transport dynamics at microtubule intersections. Proc. Natl Acad. Sci. USA 110(9):3375–3380.

51. Lukinavičius, G., L. Reymond, E. D’este, A. Masharina, F. Göttfert, H. Ta, A. Güther, M. Fournier, S. Rizzo, and H. Waldmann. 2014. Fluorogenic probes for live-cell imaging of the cytoskeleton. Nat. Methods 11(7):731.

52. Mettlen, M., P.-H. Chen, S. Srinivasan, G. Danuser, and S. L. Schmid. 2018. Regulation of clathrin-mediated endocytosis. Annu. Rev. Biochem. 87:871–896.

53. Kaksonen, M., and A. Roux. 2018. Mechanisms of clathrin-mediated endocytosis. Nat. Rev. Mol. Cell Bio 19(5):313.

54. Hohendahl, A., N. Talledge, V. Galli, P. S. Shen, F. Humbert, P. De Camilli, A. Frost, and A. Roux. 2017. Structural inhibition of dynamin-mediated membrane fission by endophilin. Elife 6:e26856.

55. Gustafsson, N., S. Culley, G. Ashdown, D. M. Owen, P. M. Pereira, and R. Henriques. 2016. Fast live-cell conventional fluorophore nanoscopy with ImageJ through super-resolution radial fluctuations. Nat. Commun. 7:12471.

56. Schaedel, L., K. John, J. Gaillard, M. V. Nachury, L. Blanchoin, and M. Théry. 2015. Microtubules self-repair in response to mechanical stress. Nat. Mater. 14(11):1156.

57. Nascimento, A. A., J. T. Roland, and V. I. Gelfand. 2003. Pigment cells: a model for the study of organelle transport. Annu. Rev. Cell Dev. Bi 19(1):469–491.

58. Lombardo, A. T., S. R. Nelson, M. Y. Ali, G. G. Kennedy, K. M. Trybus, S. Walcott, and D. M. Warshaw. 2017. Myosin Va molecular motors manoeuvre liposome cargo through suspended actin filament intersections in vitro. Nat. Commun. 8:15692.

59. Lindwall, C., and M. Kanje. 2005. Retrograde axonal transport of JNK signaling molecules influence injury induced nuclear changes in pc-Jun and ATF3 in adult rat sensory neurons. Mol. Cell Neurosci. 29(2):269–282.

60. Bronstein, I., Y. Israel, E. Kepten, S. Mai, Y. Shav-Tal, E. Barkai, and Y. Garini. 2009. Transient anomalous diffusion of telomeres in the nucleus of mammalian cells. Phys. Rev. Lett. 103(1):018102.

61. Albanese, A., P. S. Tang, and W. C. Chan. 2012. The effect of nanoparticle size, shape, and surface chemistry on biological systems. Annu. Rev. Biomed. Eng. 14:1–16.

62. Sezgin, E., I. Levental, S. Mayor, and C. Eggeling. 2017. The mystery of membrane organization: composition, regulation and roles of lipid rafts. Nat. Rev. Mol. Cell Bio 18(6):361.

63. Lemmon, M. A. 2008. Membrane recognition by phospholipid-binding domains. Nat. Rev. Mol. Cell Bio 9(2):99.

64. Antonny, B., C. Burd, P. De Camilli, E. Chen, O. Daumke, K. Faelber, M. Ford, V. A. Frolov, A. Frost, and J. E. Hinshaw. 2016. Membrane fission by dynamin: what we know and what we need to know. EMBO J. 35(21):2270–2284.

65. Lakadamyali, M., M. J. Rust, H. P. Babcock, and X. Zhuang. 2003. Visualizing infection of individual influenza viruses. Proc. Natl Acad. Sci. USA 100(16):9280–9285.

