## Supporting Information for "Resolving Cargo-motor-track Interactions in Living Cells with Bifocal Parallax Single Particle Tracking"

for

### SUPPLEMENTARY NOTES

#### 1. Parallax imaging

Parallax imaging was utilized in the current imaging setup to keep the target imaging probe in focus continuously during data collection. To achieve parallax imaging, a custom-cut wedge prism housed in a pre-optimized parallax slider was placed at the objective's back focal plane in exit pupil (Fig. S1) to create two *half-plane* images of the same sample (1). Half of the light that passed through the wedge prism was deviated by a small angle, which only used half of the numerical aperture (NA) of the objective. The other half of the light, which also utilized half of the NA of the objective, kept on the original optical path. Therefore, two half-plane images of the same imaging probe were captured in the different portions (upper and lower) on the same camera chip without overlapping. These are stated as half-plane images as they only use half of the NA of the objective.

As the pupil was evenly divided by the wedge prism, the half-plane images (upper and lower) of the imaging probe were identical for isotropic probes. Meanwhile, for anisotropic probes, such as gold nanorods (GNRs) in this study, which have transverse and longitudinal surface plasmon resonance modes (2), the split images were orientation-dependent as the two beams were divided into different profiles, depending on the orientation of GNRs relative to the wedge prism.

### 2. Effect of numerical aperture and defocusing distance on half-plane scattering imaging

Both the NA of the objective and defocusing distance affect the formation of the scattering images of GNRs. Fig. S2 shows various simulated images (full plane, half-plane upper and half-plane lower) with different orientations and defocusing distances.

Based on the characteristics of the defocusing image patterns (3), it is clear that the orientation-resolving capability is related to the NA and the defocusing distance. In dark field microscopy, the NA of the objective has to be smaller than that of the condenser (NA 1.20-1.43 in the current setup). On the other hand, the increase of the defocusing distance results in the decrease of the intensities of the patterns. These two crucial parameters must be carefully optimized to ensure good signal-to-noise ratio (S/N) of the defocused image patterns for robust pattern matching. The NA value of 1.0 and defocusing distance of 0.9 μm was chosen in this study.

### 3. Calibration of parallax imaging

In parallax imaging, when the target imaging probe is perfectly in the focal plane, the separation distance of the two half-plane mirror images is measured as $\Delta y_0$. The separation distance changes accordingly when the target probe moves along the z-axis. A set of half-plane images was obtained from a z-scan of the sample.

To generate a calibration curve of $\Delta y$ as a function of $\Delta z$, over 30 GNRs immobilized on a glass slide surface with different orientations were scanned along z-axis from -1000 nm to 1000 nm with 10 nm scanning step using a high-precision objective scanner (Fig. S3). The accurate positions of the target GNRs were obtained to establish the calibration curve between $\Delta y$ and $\Delta z$.

Based on the calibration curve, an auto-feedback z adjustment system was implemented with a closed-loop nanometer-precision piezo stage and a tracking program in ImageJ μManger (4, 5). When the automatic feedback loop system is enabled, the separation distance will be obtained in every frame. If the separation distance is larger or smaller than $\Delta y_0$, the z piezo stage will move to a certain value based on the calibration curve within a response time of less than 1 ms. More importantly, the z coordinate of target was provided by the z piezo stage physical output system which ensures the high accuracy.

### 4. 3D localization precision

The ellipse-shaped half-plane scattering images in the focused channel can be fitted by 2D elliptical Gaussian functions for determining the GNR position:

$$I(x,y) = A + B * exp(-(\frac{(x-x_0)^2}{2S_x^2} + \frac{(y-y_0)^2}{2S_y^2})) \quad \text{(S1)}$$

where $(x_0, y_0)$ is the center position, A is the background level, B is the peak intensity at $(x_0, y_0)$, $S_x$ and $S_y$ are the standard deviations of the Gaussian distribution along x- and y-axis, respectively.

Several factors such as photon noise, scattering background, camera readout noise, and pixel size can influence the half-plane scattering images captured by the EMCCD camera. The localization precision ($\sigma_{j\ ,j=x,\,y}$) can be calculated according to equation (S2) (6, 7):

$$\sigma_j = \sqrt{\left(\frac{S_j^2}{N} + \frac{a^2/12}{N} + \frac{8\pi S_j^4 b^2}{a^2 N^2}\right)} \quad \text{(S2)}$$

where $N$ is the photons collected in each imaging frame, $a$ is the pixel size, and $b$ is the background noise in photons. The x and y coordinates can be resolved by correlation mapping in the focused channel, and the z coordinate can be obtained from the recorded movement of the objective scanner. In the specific example in Fig. S4 (left), the localization precisions are determined to be $\sigma_x$ = 2.1 nm, $\sigma_y$ = 4.8 nm for the particle in upper panel given the parameters of $S_x$ = 138 nm, $S_y$ = 235 nm, $N$ = 6914 photons, $b$ = 13 photons, and the pixel size of image $a$ = 131 nm; the localization precisions are determined to be $\sigma_x$ = 2.3 nm, $\sigma_y$ = 6.1 nm for the particle in lower panel given the parameters of $S_x$ = 139 nm, $S_y$ = 266 nm, $N$ = 6366 photons, $b$ = 12 photons. In the current setup for determining the GNRs position in z-axis, we use $z = k(y_{upper} - y_{lower})$, where $k$ is the slope of the calibration curve between z and Δy (Fig. S3). The localization precision $\sigma_z$ is calculated from the propagation of error:

$$\sigma_z = k\sqrt{\sigma^2{}_{y_{upper}} + \sigma^2_{y_{lower}} - 2\sigma_{y_{upper}y_{lower}}} \quad \text{(S3)}$$

where $\sigma_{y_{upper}y_{lower}}$= 0 for independent measurements of the upper and lower half-plane images. Specifically, the localization precision $\sigma_z$ for the example in Fig. S4 (left) is determined to be 11.4 nm.

We collected imaging data for 1000 GNRs and determined their positions using the same method as discussed above. The localization precisions in 3D are determined statistically (Fig. S4), giving $\sigma_x = 2.2 \pm 0.1$ nm, $\sigma_y = 4.8 \pm 0.3$ nm in upper channel, $\sigma_x = 2.4 \pm 0.1$ nm, $\sigma_y = 6.4 \pm 0.4$ nm in lower channel, and $\sigma_z = 11.6 \pm 0.5$ nm. The localization precision is worse in y compared to that in x because the half-plane point spread functions (PSFs) are stretched in y in our imaging setup. Since the localization precisions is better in the upper channel in our imaging setup, we used the determined center positions in the upper channel for 2D positions of GNRs in our experiments.

### 5. Simulation of half-plane scattering point spread functions

Computer simulation of PSF is essential in acquiring a complete understanding of the experimental image patterns and to estimate uncertainties in the related measurements. Several methods have been developed to determine the dipole orientation by fitting experimental images with the simulated patterns.(8, 9) However, none of the existing simulation programs can be used directly for parallax microscopy, which employs a semi-circle pupil to result in the half-plane PSFs. To fully understand the irregular shapes of the defocused half-plane images, a simulation program to obtain theoretical half-plane PSFs as a function of azimuth and polar angles using the six basic functions of dipole emission ($I_x^2$ , $I_{xy}$ , $I_y^2$ , $I_{xz}$ , $I_z^2$ , $I_{yz}$ ) (10, 11) has been developed in house .

In the dark field microscopy, the GNRs can be illuminated by unpolarized light through an oil-immersion dark field condenser and the scattering images are collected by a NA 1.0 objective. The anisotropic object can be defined as a multi-oscillation dipole, and the far-field intensity distribution of a dipole is characterized by a spatial position $x_{ff} = (x_{ff}, y_{ff}, z_{ff})$ and orientation angle which can be described as a unit vector $\hat{r}$ or an azimuth angle $\theta$ and polar angle $\phi$, where $\hat{r} = (\sin\theta_{ff}\cos\phi_{ff}, \sin\theta_{ff}\sin\phi_{ff}, \cos\theta_{ff})$. The measured scattering spot can be written as (9):

$$
\begin{aligned}
h_{\theta_{ff},\phi_{ff}}\left(x; x_{ff}, \tau\right) &= |\varepsilon|^2 \\
&= \sin^2\theta_{ff}\left(|I_0|^2 + |I_2|^2 + 2\cos\left(2\phi_{ff} - 2\phi_d\right)\Re\left\{I_0^* I_2\right\}\right) \\
&\quad -2\sin(2\theta_{ff})\cos(\phi_{ff} - \phi_d)\Im\left\{I_0 + I_2\right\} + 4|I_1|^2\cos_2\theta_{ff} \\
&= \hat{r}^T M_{ff}
\end{aligned}
$$

(S4)

where $\phi_d = \tan^{-1}((y - y_{ff})/(x - x_{ff}))$. An asterisk represents the adjoint operator. $\hat{r}^T$ denotes the conjugate transpose of $\hat{r}$, $\Re\{v\}$ and $\Im\{v\}$ stand for the real and imaginary components of $v$, and $\tau$ is the parameters of the optical setup. $M_{ff}$ is a symmetric matrix which contains six non-orthogonal templates given by:

$$\begin{aligned}
m_{11} &= |I_0|^2 + |I_2|^2 + 2\Re\{I_0^* I_2\}\cos 2\phi_d \\
m_{12} &= 2\Re\{I_0^* I_2\}\sin 2\phi_d \\
m_{13} &= -2\cos\phi_d \Im\{I_1^*(I_0 + I_2)\} \\
m_{22} &= |I_0|^2 + |I_2|^2 - 2\Re\{I_0^* I_2\}\cos 2\phi_d \\
m_{23} &= -2\sin\phi_d \Im\{I_1^*(I_0 + I_2)\} \\
m_{33} &= 4|I_1|^2
\end{aligned} \tag{S5}$$

where

$$I_0(x; x_{ff}, \tau) = \int_0^\alpha B_0(\theta)\left(t_s^{(1)} t_s^{(2)} + t_{ff}^{(1)} t_{ff}^{(2)} \frac{1}{n_s}\sqrt{n_s^2 - n_i^2 \sin^2\theta}\right) d\theta$$

$$I_1(x; x_{ff}, \tau) = \int_0^\alpha B_1(\theta) t_{ff}^{(1)} t_{ff}^{(2)} \frac{n_i}{n_s} \sin\theta d\theta \tag{S6}$$

$$I_2(x; x_{ff}, \tau) = \int_0^\alpha B_2(\theta)\left(t_s^{(1)} t_s^{(2)} - t_{ff}^{(1)} t_{ff}^{(2)} \frac{1}{n_s}\sqrt{n_s^2 - n_i^2 \sin^2\theta}\right) d\theta$$

with
$$B_m(\theta) = \sqrt{\cos\theta}\sin\theta J_m(krn_i \sin\theta) e^{ik\Lambda(\theta, z, z_{ff}, \tau)} \tag{S7}$$

where the integrals for $I_0$, $I_1$ and $I_2$ are the vectoral PSF equation. As the $\Lambda(\theta, z, z_{ff}, \tau)$ denotes the system's aberrations affected by defocusing.

In our optical setup, as the wedge prism was placed at the back focal plane of the objective, only half of the pupil was used for the formation of each half-plane image. Therefore, equation (S1) can be rewritten as:

$$h_{\theta_{ff},\phi_{ff}}\left(x;x_{ff},\tau\right)=\hat{r}^{T}\,[P_{ff}M_{ff}] \tag{S8}$$

where $P_{ff}$ represents a pupil function. The six basic full-plane dipole emission templates are shown in Fig. S6. By introducing a pupil function $P_{ff}$, two sets of six basis templates for the upper and lower half-plane PSFs have also been simulated and shown in Fig. S6. Using a linear combination of these basis templates, theoretical PSFs (the focused upper and lower, and defocused upper and lower PSFs) as a function of dipole orientation can be simulated. Compared to the previous methods that extracted a simplified function between orientation (polar angle and azimuth angle) and scattering intensities by omitting higher-order polynomials, our method is advantageous by retaining all polynomials to decrease the uncertainty of angular determination in pattern matching with the experimental images.

These basic patterns are dependent on system-specific parameters, including the numerical aperture, magnification of the objective, and defocusing distance, and their linear combination can reproduce the half-plane image patterns.

### 6. Uncertainties of angle determination

The uncertainties associated with the azimuth and polar angles are orientation dependent. Under typical live cell imaging conditions, the S/N ratio of 50 is readily attainable for regions with relatively low background. Simulated images with artificial noise (S/N = 50) were used as the calibration data set (with 1° interval for both polar and azimuth angles, 50 images per position). When the polar angle is between 10-80°, a precision of <1° can be recovered for any combination of the polar angle and the azimuth angle (Fig. S7 and S8 for azimuth angle and polar stepping, respectively, image S/N = 50). When the polar angle is very small (i.e., a standing nanorod), the image patterns become similar to each other and the azimuth angle can no longer be recovered (random φ being reported). When the polar angle is very close to 90° (i.e., a nanorod lying flat), the azimuth angles become degenerated and can only be recovered for the range of 0-180°, which is expected because the asymmetry in the PSF disappears when the nanorod lies flat on a horizontal plane.

When the S/N is reduced to ~10 in cellular regions with relatively high background, both the polar angle and the azimuth angles can still be recovered with a precision of < 2° for the polar angle range of 10-80°.

**Table S1. Parameters describing the translation and rotational motions of GNRs.**

| | α | Diffusion coefficient (μm$^2$/s) | Step azimuth angle (°) | Step polar angle (°) |
|---|---|---|---|---|
| Free diffusion | 0.98± 0.20 | 0.22± 0.14 | 72.4± 84.5 | 17.4 ± 21.0 |
| Active transport | 1.68± 0.28 | 0.08± 0.15* | 4.9 ± 7.0 | 4.0 ± 5.5 |
| Tight attachment | 0.49±0.28 | 0.0017± 0.001 | 4.9± 6.7 | 4.3 ± 5.7 |
| Tethered rotation | 0.50±0.29 | 0.004± 0.004 | 12.5± 22.5 | 7.0 ± 9.7 |

Note: Data are presented as Mean ± Standard Deviation unless otherwise noted. The event number (n) for each data point of α and diffusion coefficient is >50. For the step azimuth and polar angle, each data point >1500.

* For the active transport cases, the linear velocity is 1.2 ± 1.0 μm/s.

### SUPPLEMENTARY FIGURES

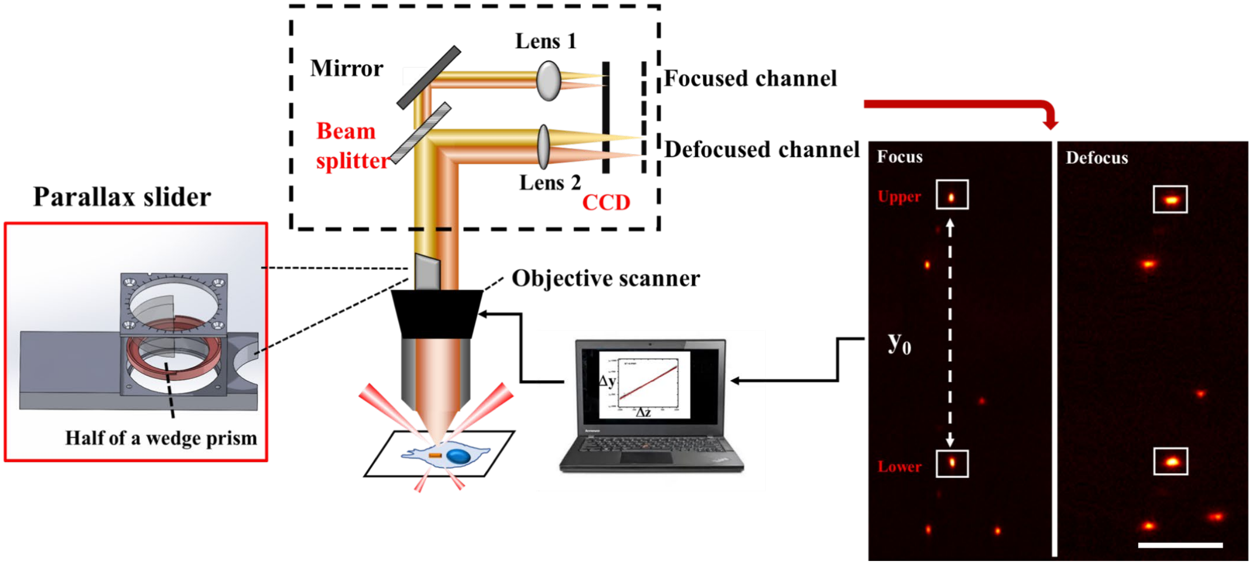

**Fig. S1.** Schematics of bifocal parallax SPT setup. The black dashed line square indicates the bifocal imaging system. Four spots (upper and lower in-focus spots and upper and lower defocus patterns) were generated by the imaging system for a single GNR. The upper and lower in-focus spots have the same x coordinate and the distance between them is $y_0$ for a GNR perfectly in the focal plane. The defocusing distance in the defocus channel is optimized to 0.9 μm. Scale bar is 10 μm.

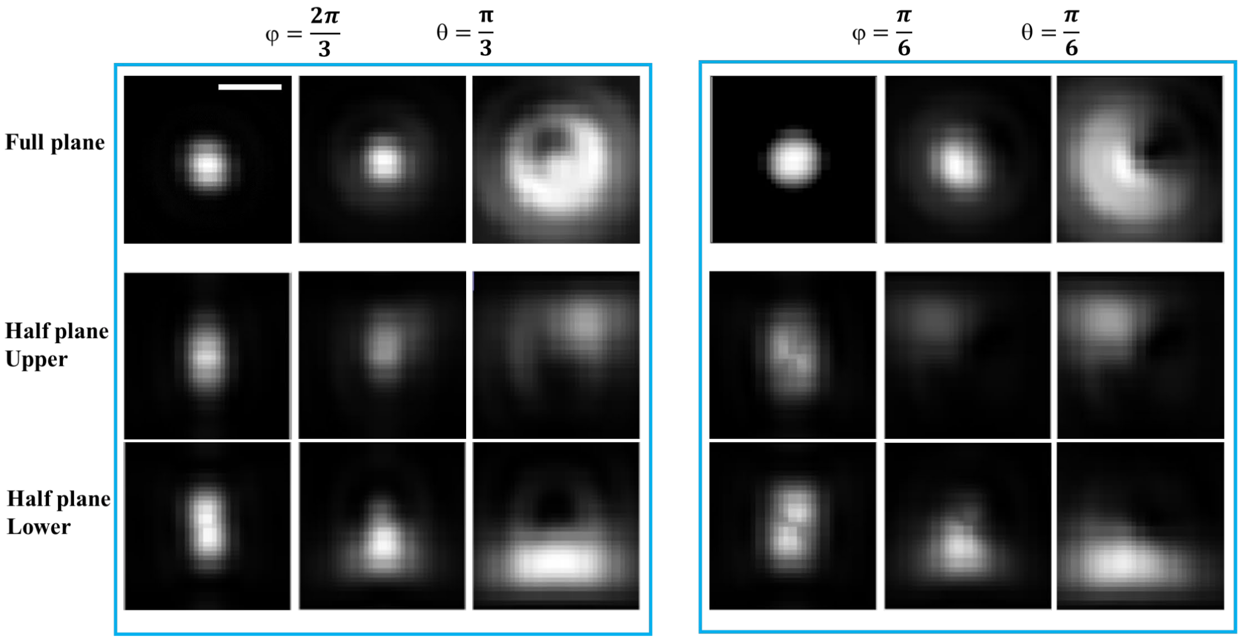

**Fig. S2.** Simulated full-plane and half-plane scattering PSFs at various orientations and defocusing distances. Scale bar is 1 µm. In both light blue boxes, the defocusing distance is 0 µm (left column), 0.5 µm (middle column), 0.9 µm (right column).

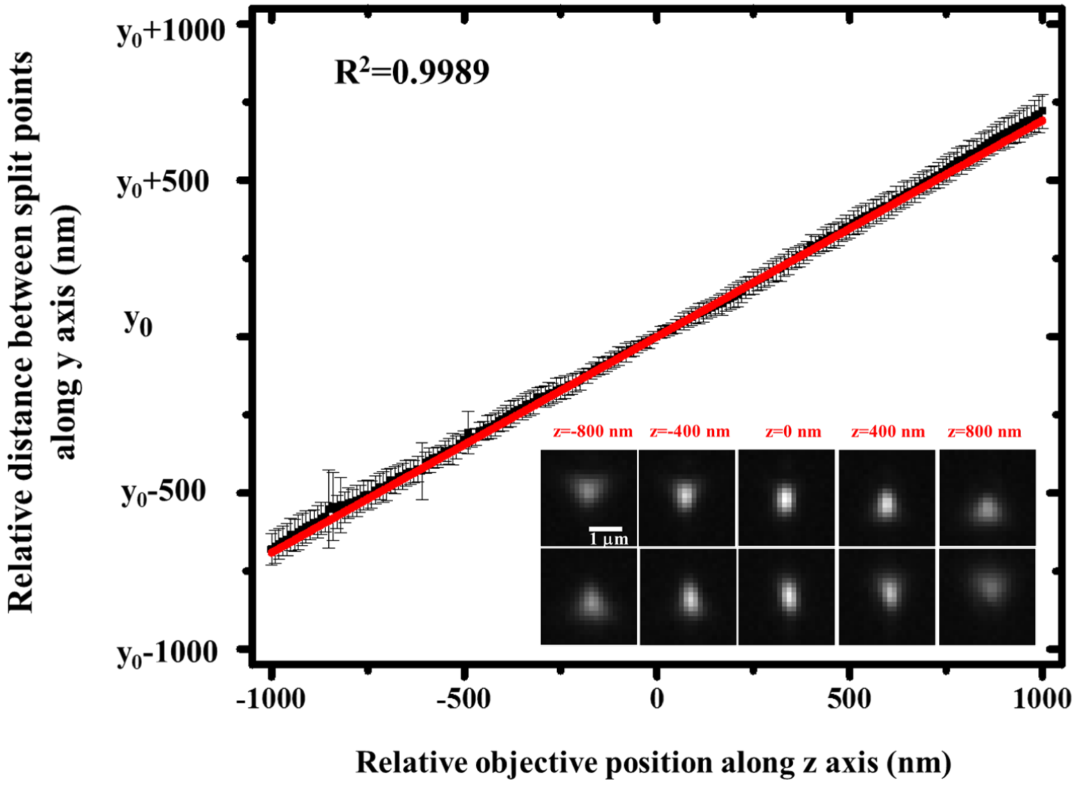

**Fig. S3.** Calibration curve of Δy *vs.* Δz. Over 30 GNRs immobilized on a glass slide surface with various orientations were scanned along the z-axis from -1000 nm to 1000 nm with 10 nm step using the high-precision objective scanner. The inserted images are pairs of half-plane scattering imaging (upper and lower) at various vertical positions.

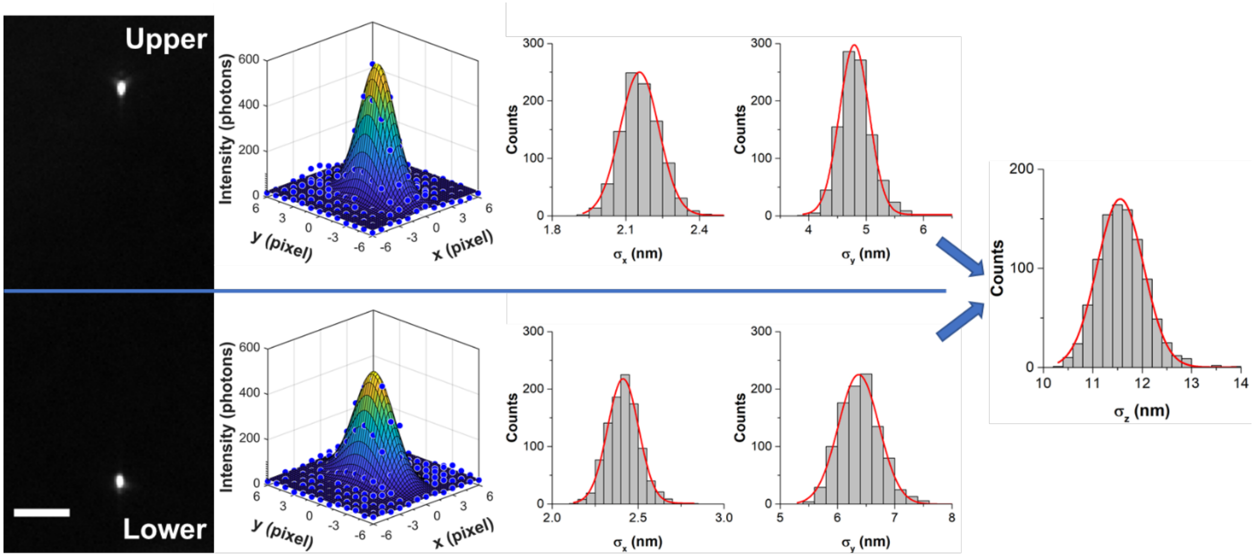

**Fig. S4.** 3D localization of GNRs. Typical upper and lower half-plane dark-field images of a GNR with 0.02 s integration time in the focused channel are shown to the left. The center positions are determined by Gaussian fitting and localization precisions are determined to be $\sigma_x$ = 2.1 nm, $\sigma_y$ = 4.8 nm for the particle in the upper panel and $\sigma_x$ = 2.3 nm, $\sigma_y$ = 6.1 nm for the particle in the lower panel. Gaussian fitting the histogram distributions of localization precisions over 1000 single GNRs gives the statistical values of $\sigma_x$ = 2.2 ± 0.1 nm, $\sigma_y$ = 4.8 ± 0.3 nm (upper) and $\sigma_x$ = 2.4 ± 0.1 nm, $\sigma_y$ = 6.4 ± 0.4 nm (lower). The localization precision in z is calculated to be $\sigma_z$ = 11.6 ± 0.5 nm using the error propagation method (***SI text***). The average localization precisions are presented as Mean ± SD from the Gaussian fitting the histogram distributions of localization precisions. Scale bar is 5 μm.

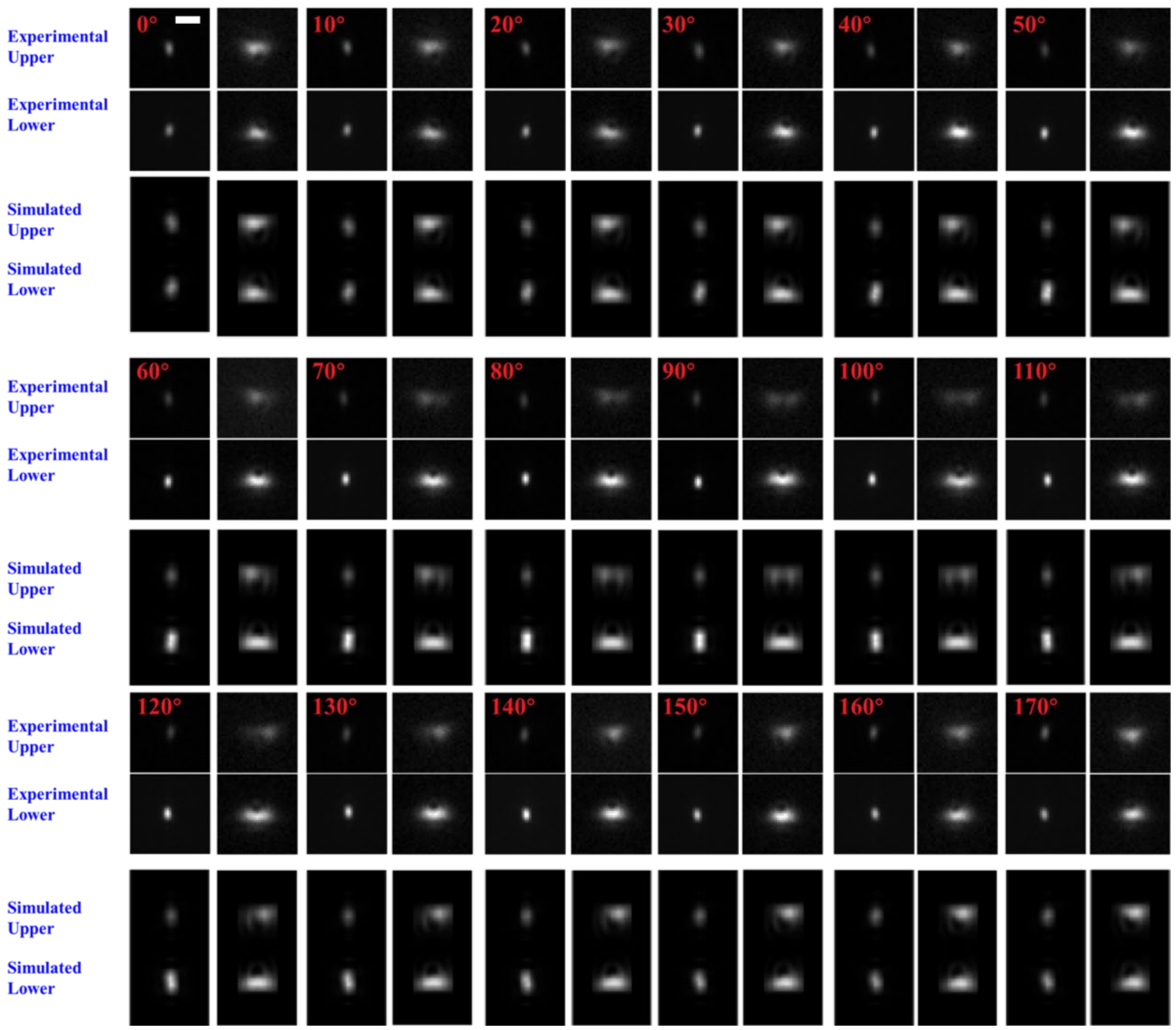
Experimental Upper
Experimental Lower
Simulated Upper
Simulated Lower
0°
10°
20°
30°
40°
50°
60°
70°
80°
90°
100°
110°
120°
130°
140°
150°
160°
170°

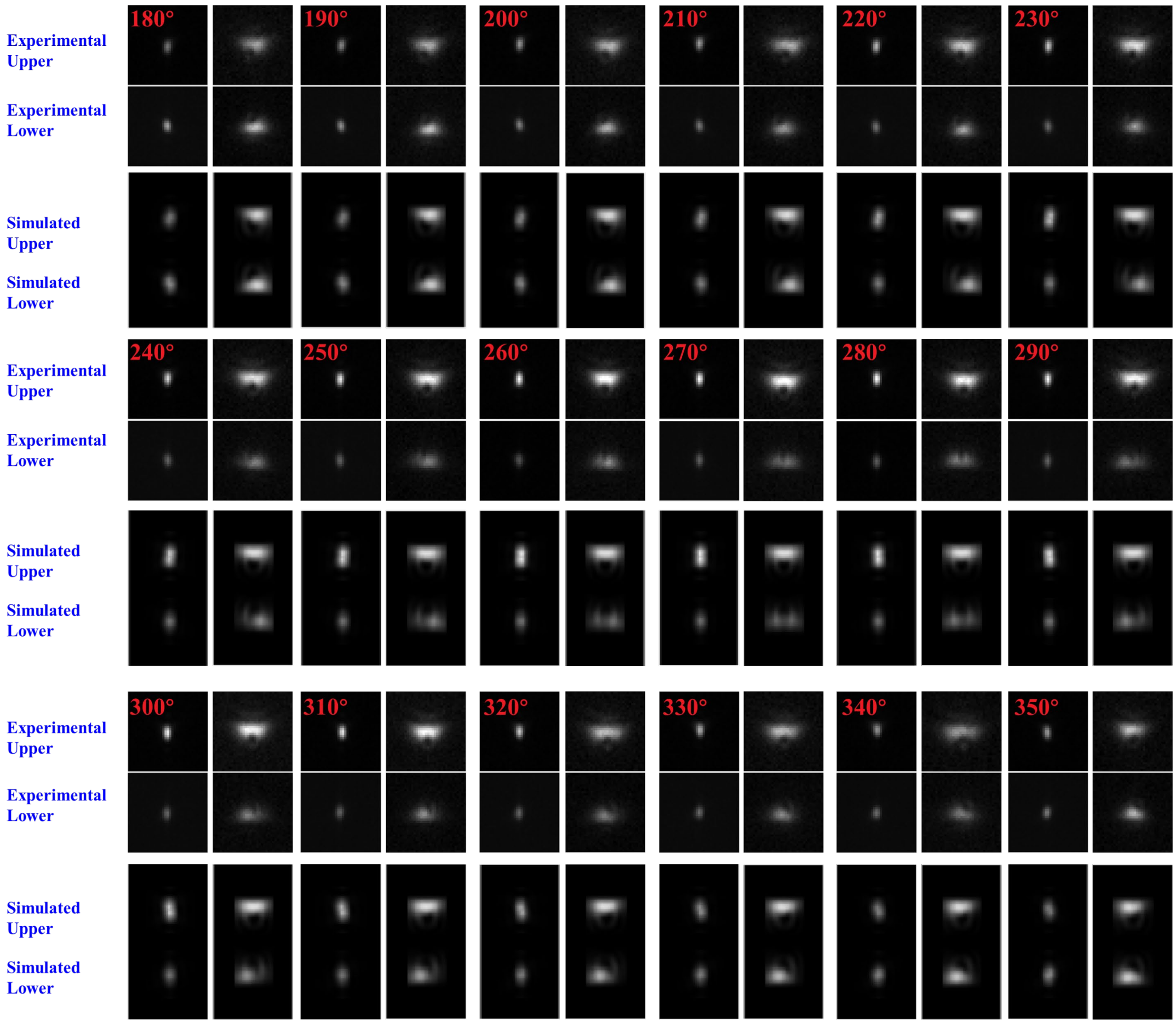

**Fig. S5.** Experimental and simulated half-plane scattering images (the focused upper and lower images on the left and the defocused upper and lower images on the right) of a GNR with a polar angle of 60° as a function of azimuth angle with a 10° interval. Scale bar is 2 µm.

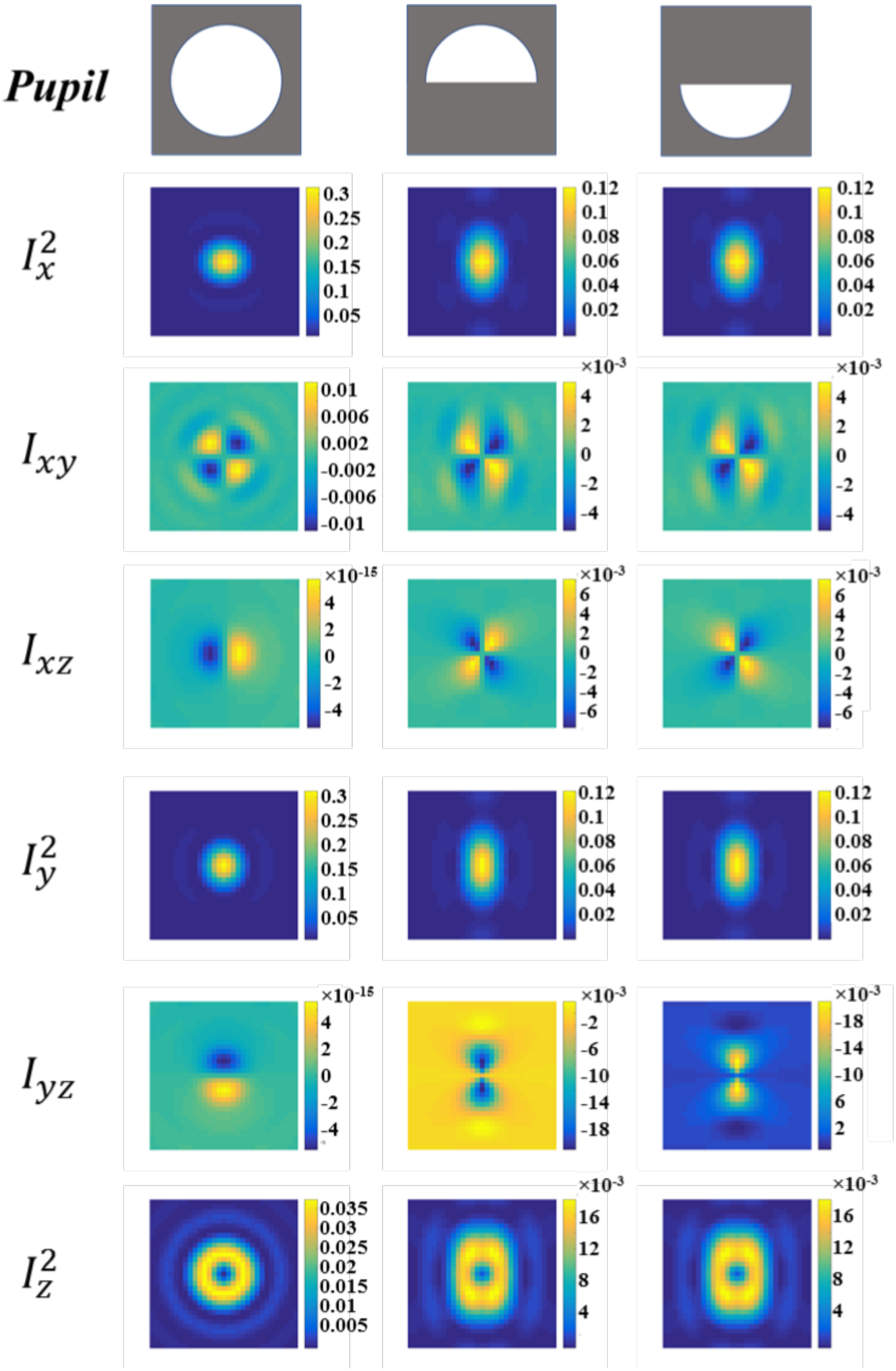

**Fig. S6.** Simulated pupil and six basis dipole emission templates: full-plane (left column), half-plane upper spot (middle column), half-plane lower spot (right column).

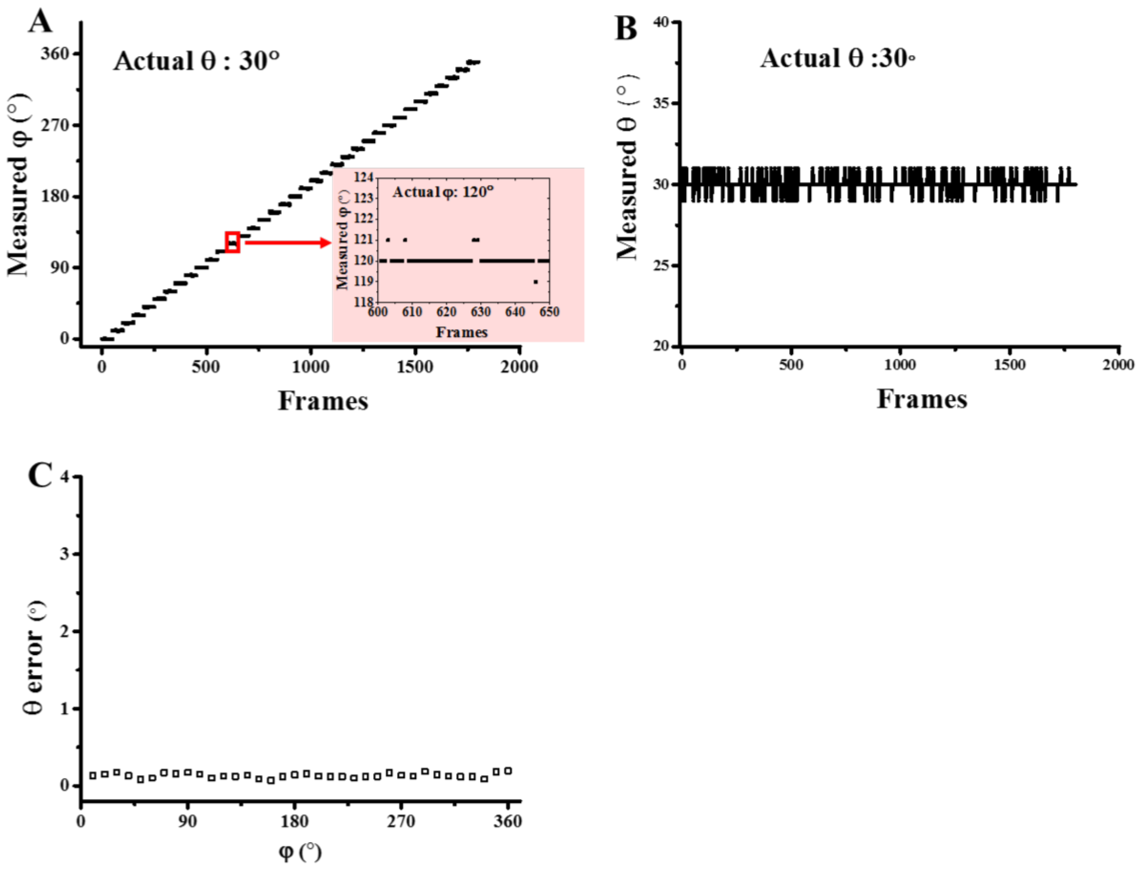

**Fig. S7.** Stepping experiment demonstrating the precision of pattern recognition at S/N of 50. (A, B) Polar angle (θ) was set at 30° and azimuth angle (φ) was stepped at 10°, and 50 images were taken for each step. (A) is the recovered azimuth angle and (B) is the recovered polar angle values. The inserted graph showed the higher-magnification views of the red square regions in which the actual azimuth angle and polar angle was 120° and 30°, respectively. (C) Estimated standard errors for the polar angle with various azimuth angles.

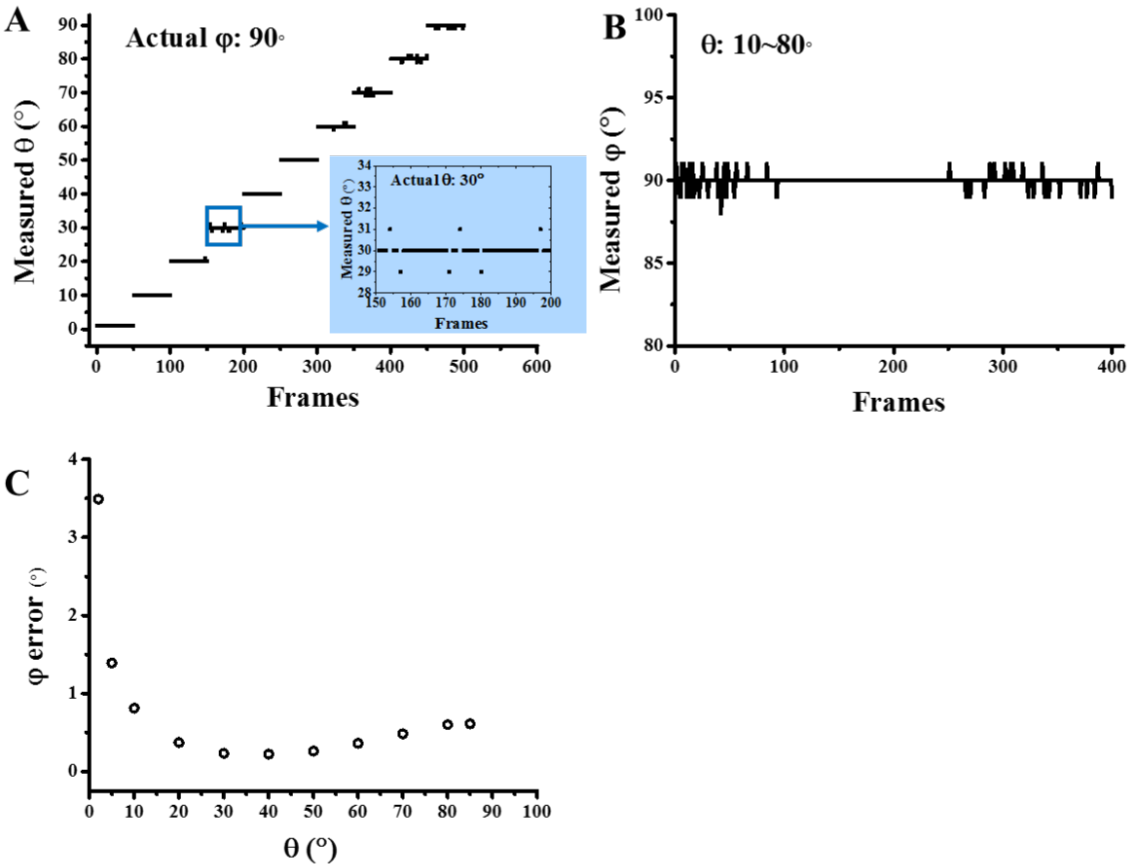

**Fig. S8.** Stepping experiment demonstrating the precision of pattern recognition at S/N of 50. (A, B) azimuth angle (φ) was set at 90° and polar angle (θ) was stepped at 10°, and 50 images were taken for each step. (A) The recovered azimuth angle and (B) The recovered polar angle values, respectively. The inserted graph showed the higher-magnification views of the blue square regions in which the actual azimuth angle and polar angle was 90 and 30°, respectively. (C) Estimated standard errors for azimuth angle with various polar angles.

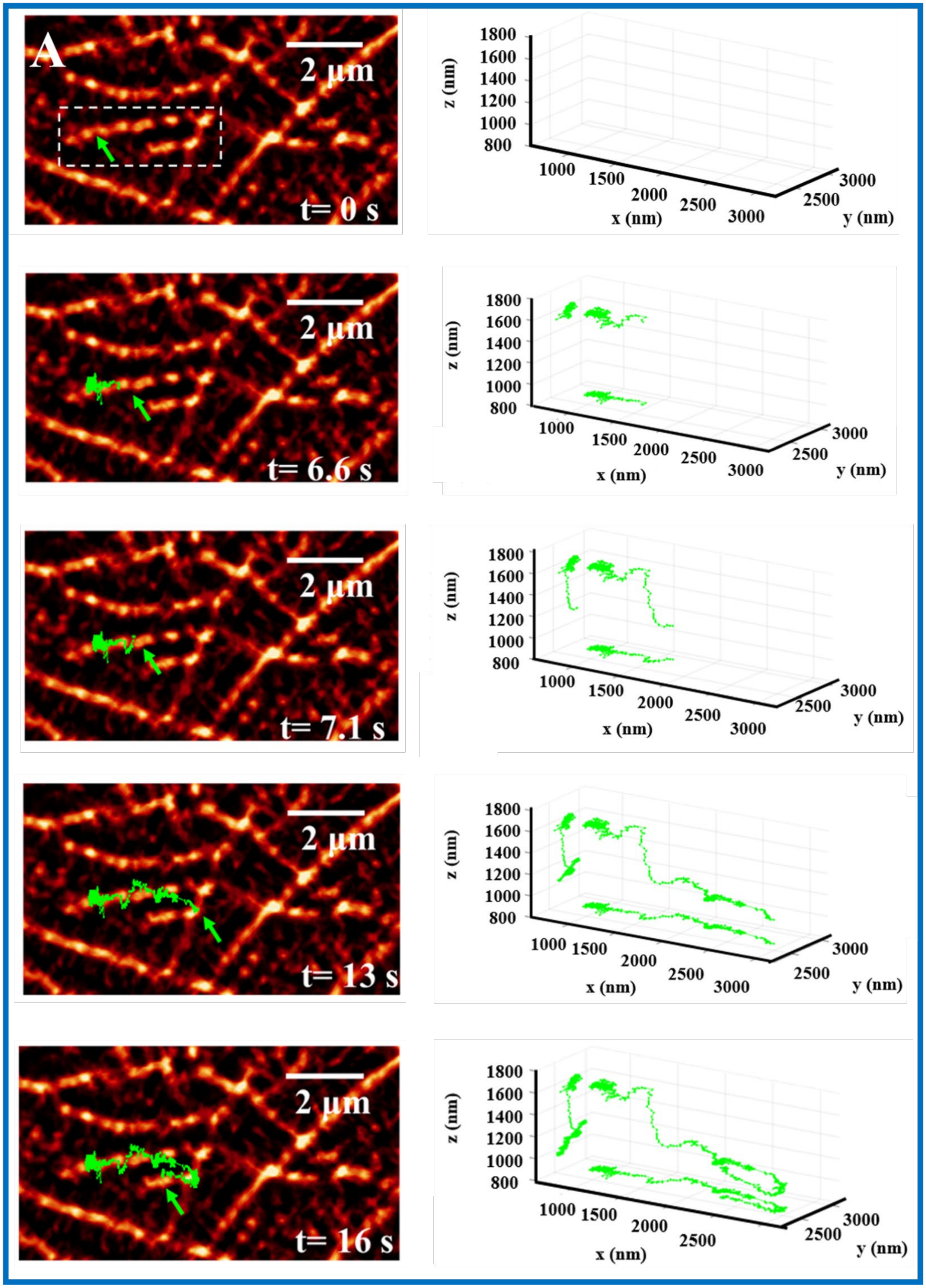
A
2 µm
t= 0 s
z (nm)
1800
1600
1400
1200
1000
800
1000
1500
2000
2500
3000
x (nm)
3000
2500
y (nm)
t= 6.6 s
t= 7.1 s
t= 13 s
t= 16 s

B

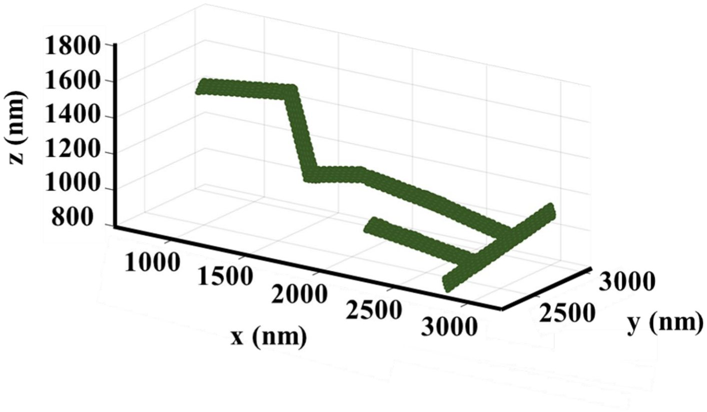

**Fig. S9.** (A) Overlap of live-cell imaging trajectory of a cargo and the SRRF microtubule images (Left) and corresponding 3D trajectory of a cargo with projection on xy- plane and xz- plane (Right) at various time ranges. The green arrow shows the endpoint of the trajectory of the cargo. (B) The resolved 3D microtubule structure of the white dashed square in (A).

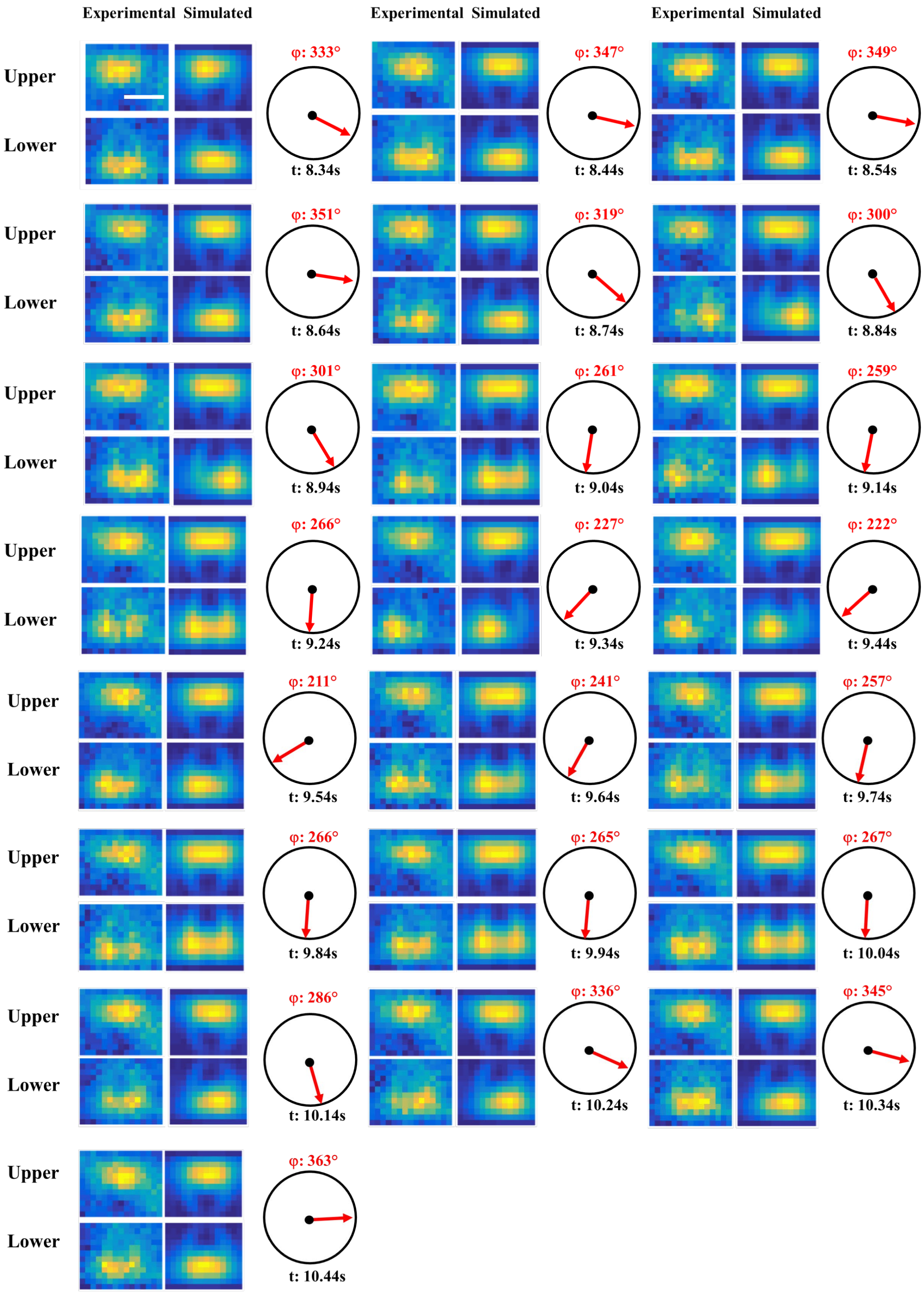
Experimental Simulated
Experimental Simulated
Experimental Simulated
Upper
Lower
φ: 333°
t: 8.34s
φ: 347°
t: 8.44s
φ: 349°
t: 8.54s
φ: 351°
t: 8.64s
φ: 319°
t: 8.74s
φ: 300°
t: 8.84s
φ: 301°
t: 8.94s
φ: 261°
t: 9.04s
φ: 259°
t: 9.14s
φ: 266°
t: 9.24s
φ: 227°
t: 9.34s
φ: 222°
t: 9.44s
φ: 211°
t: 9.54s
φ: 241°
t: 9.64s
φ: 257°
t: 9.74s
φ: 266°
t: 9.84s
φ: 265°
t: 9.94s
φ: 267°
t: 10.04s
φ: 286°
t: 10.14s
φ: 336°
t: 10.24s
φ: 345°
t: 10.34s
φ: 363°
t: 10.44s

**Fig. S10.** Experimental and simulated half-plane scattering images (upper and lower in-focus spots, upper and lower defocus patterns) of the cargo in the final period (8.34-10.5s) of directional rotation in every 5 frames (a time interval of 100ms) of Fig. 4. The inserted cartoons indicate the direction of azimuth angle of cargo, and the inserted φ and t show the values of azimuth angle of cargo and time. Scale bar is 1 μm. The result shows the cargo experienced gradual azimuth angle change (a 150° counterclockwise rotation followed by another 150° clockwise rotation).

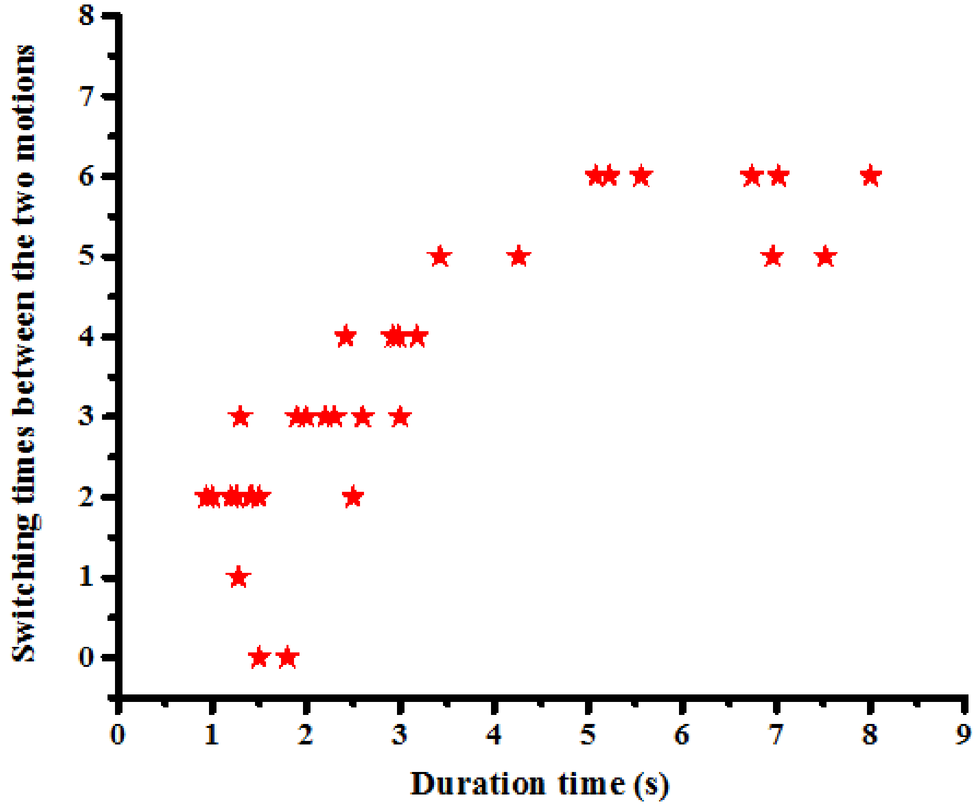

**Fig. S11.** The distribution of the pause duration with number of switches between tight attachment and tethered rotation.

### SUPPLEMENTARY MOVIE

All movies were captured at 50 fps and played at a slower speed of 20 fps.

**Movie S1.** Corresponding movie of Fig. 3. A cargo moves across a U-shape microtubule intersection.

**Movie S2.** 3D reconstitution of Movie S1. The inserted orange rod indicates the GNR cargo. The inserted t, theta, phi and z show the time, azimuth angle, polar angle and z coordination of the cargo.

**Movie S3.** Corresponding 3D reconstitution movie of Fig. 4. A cargo moves across a microtubule intersection with switching between tight attachment and tethered rotation. The inserted orange rod and symbol represent the same meanings with those of Movie 2.

### SUPPLEMENTARY REFERENCES

1. Chen, K., Y. Gu, W. Sun, G. Wang, X. Fan, T. Xia, and N. Fang. 2017. Characteristic rotational behaviors of rod-shaped cargo revealed by automated five-dimensional single particle tracking. Nat. Commun. 8(1):887.
2. Huang, X., I. H. El-Sayed, W. Qian, and M. A. El-Sayed. 2006. Cancer cell imaging and photothermal therapy in the near-infrared region by using gold nanorods. J. Am. Chem. Soc. 128(6):2115-2120.
3. Xiao, L., Y. Qiao, Y. He, and E. S. Yeung. 2010. Three dimensional orientational imaging of nanoparticles with darkfield microscopy. Anal. Chem. 82(12):5268-5274.
4. Schneider, C. A., W. S. Rasband, and K. W. Eliceiri. 2012. NIH Image to ImageJ: 25 years of image analysis. Nat. Methods 9(7):671.
5. Edelstein, A., N. Amodaj, K. Hoover, R. Vale, and N. Stuurman. 2010. Computer control of microscopes using µManager. Curr. Protoc. Mol. Biol. 92(1):14.20. 11-14.20. 17.
6. Yildiz, A., J. N. Forkey, S. A. McKinney, T. Ha, Y. E. Goldman, and P. R. Selvin. 2003. Myosin V walks hand-over-hand: single fluorophore imaging with 1.5-nm localization. Science 300(5628):2061-2065.
7. Thompson, R. E., D. R. Larson, and W. W. Webb. 2002. Precise nanometer localization analysis for individual fluorescent probes. Biophys. J. 82(5):2775-2783.
8. Patra, D., I. Gregor, and J. Enderlein. 2004. Image analysis of defocused single-molecule images for three-dimensional molecule orientation studies. J. Phys. Chem. A 108(33):6836-6841.
9. Aguet, F., S. Geissbühler, I. Märki, T. Lasser, and M. Unser. 2009. Super-resolution orientation estimation and localization of fluorescent dipoles using 3-D steerable filters. Opt. Express 17(8):6829-6848.
10. Burghardt, T. P. 2011. Single molecule fluorescence image patterns linked to dipole orientation and axial position: application to myosin cross-bridges in muscle fibers. PLoS One 6(2):e16772.
11. Backer, A. S., and W. Moerner. 2014. Extending single-molecule microscopy using optical Fourier processing. J. Phys. Chem. B 118(28):8313-8329.
